# Proteome Reallocation Reveals Dynamics and Mechanisms of Phage Infection and Defense in *Staphylococcus aureus*

**DOI:** 10.1101/2025.10.23.684294

**Authors:** David Dooley, Dana L. Carper, Richard J. Giannone, Cong T. Trinh

## Abstract

The coexistence of bacteria and phages is marked by dynamic interactions that determine infection outcomes. However, the mechanisms by which the host allocates its resources to cope with phage infection remain largely unknown. In this study, using longitudinal proteomics, we elucidated these interactions for the temperate staphylococcal phage *ϕ*NM1 and strains of *Staphylococcus aureus* either harboring its cognate prophage or lacking it. We demonstrated that infection of non-lysogenic *S. aureus* with *ϕ*NM1 induces a dramatic shutdown of host translation, reducing proteome allocation by over 20%. Quantitative analysis of the economics of *ϕ*NM1 infection revealed that the diversion of these cellular resources toward phage replication imposes a significant metabolic burden, thereby impairing cell growth. In contrast, lysogenic cells cope with phage infection and prevent culture collapse through a coordinated response of prophage-encoded defenses, host-encoded stress effectors, and reprogrammed cellular metabolism, thereby avoiding translation shutdown. Through coinfection with the wildtype phage and an engineered phage-like particle carrying a CRISPR-Cas phagemid, we revealed that synthetic DNA cargos evade host defenses and hijack the transcriptional machinery, altering infection outcomes. Without immunity, coinfection could collapse the non-lysogens more quickly than the native phage by overexpressing the cargo proteins, suppressing carbohydrate metabolism, and accelerating structural phage protein production through increased phage genome replication. Together, these findings provide a systems-level understanding of phage infection in *S. aureus*, uncovering the mechanisms for host takeover and prophage-mediated defense, with implications for next-generation phage therapy.

**Significance:** Multidrug-resistant bacteria pose a major threat to human health. Phage therapy offers a precise therapeutic approach by leveraging the specificity of phage infection and cargo delivery. However, as with conventional antibiotics, phage resistance can develop. Understanding the dynamic interactions between bacterial hosts and phages is therefore essential for predicting infection outcomes and designing precision phage therapies to suppress resistance. Using longitudinal proteomics, we elucidated the dynamics by which bacterial hosts reallocate cellular resources to cope with phage infection at the systems level. Coordinated defenses and reprogrammed metabolism are critical for host survival, but phage-delivered cargos can effectively bypass these barriers. These insights into the intricate interplay between the phage and host are crucial for designing next-generation precision phage therapies with greater lethality and less resistance.

## Introduction

Phages have historically been viewed as mortal enemies of bacteria, but current understanding challenges this notion. Phages exist in either a lytic cycle or a lysogenic cycle(1). A lytic cycle is characterized by acute infection, replication, and propagation of phage progeny via bacterial lysis, while a lysogenic cycle involves quiescent replication of the prophage on a plasmid, or more commonly, integrated within the host chromosome(2). In the latter case, prophages frequently confer increased fitness and/or virulence to the host in a phenomenon known as lysogenic conversion(3-5). Therefore, phages can be beneficial, harmful, or neutral to bacterial hosts depending on environmental and genetic factors.

*Staphylococcus aureus* has emerged as a leading threat to human health due to its clinical severity and aggressive acquisition of antimicrobial resistance(6-8). In response, phage therapy has resurfaced as a promising alternative to traditional antibiotics, which are dwindling in their efficacy(9, 10). Several studies have demonstrated the potential of natural and engineered phage-based therapies for combating virulent *S. aureus*(11-19), but a fundamental understanding of their modes of action and vulnerabilities remains limited. It is well-known that phages ultimately lyse infected cells through the combined activity of holin and endolysin, but before this can occur, phages must overcome bacterial immune responses, make lysis or lysogeny decisions(20), hijack host cell machinery(21, 22), and assemble nascent virions. The interplay between these processes and host metabolic and stress responses remains underexplored, especially in this clinically relevant pathogen.

Phage therapy faces the additional challenge of superinfection exclusion, whereby prophages provide their hosts with immunity to infection by similar phages. Since approximately 70–90% of *S. aureus* isolates possess at least one prophage(23-26), with most possessing more than one, understanding how prophages defend against phage infection and manipulate host metabolism is paramount for developing effective next-generation phage therapies. One strategy to overcome prophage defenses is to include an orthogonal antimicrobial payload within the therapeutic phage, either on a phagemid or within the phage genome itself(27). Therapeutic phagemids typically express several proteins like selection markers, replication machinery, and/or antimicrobial effectors (e.g., CRISPR-Cas), which can significantly alter cellular function and phage-host interactions, but these effects are not well-understood, highlighting the need for systems-level interrogation.

To address these knowledge gaps, we employed physiological characterization and longitudinal proteomics to quantitatively evaluate the effects of staphylococcal phage *ϕ*NM1 infection on the cell growth kinetics, phage proliferation, proteomic response, and resource allocation of the prophage-deficient *S. aureus* RN4220 (non-lysogen) and its *ϕ*NM1 lysogen, RN4220*^ϕ^*^NM1^ (SaDD0001, lysogen). To elucidate the effect of phage-borne DNA cargos on infection outcomes in non-lysogenic and lysogenic *S. aureus*, we reprogrammed *ϕ*NM1 to create phage-like particles that use *ϕ*NM1 capsids to package a null-targeting CRISPR-Cas phagemid and investigated their coinfection with *ϕ*NM1.

## Results and Discussion

### Lysogenic and non-lysogenic *S. aureus* exhibited distinct growth and altered proteomes in response to *ϕ*NM1 infection

To investigate the dynamic interplay between the host and phage during infection, we inoculated exponentially grown *S. aureus* strains, RN4220 and SaDD0001, at an initial OD of 0.10 and infected them with the phage *ϕ*NM1 at a multiplicity of infection (MOI) of 1 when the cells reached an OD of 0.6–0.9 (**Fig. 1A**). Samples from the non-lysogen and lysogen cultures were collected at pre-, mid-, and late-infection time points in response to *ϕ*NM1, and proteomes were measured alongside cell growth kinetics. Even though both the non-lysogen (0.524±0.052 h^-1^) and lysogen (0.541±0.021 h^-1^) grew at similar maximum specific growth rates during pre-infection, they exhibited distinct phenotypes following infection (**Fig. 1B**). The non-lysogen showed moderate growth inhibition (0.373±0.010 h^-1^) during mid-infection and severe growth inhibition during late-infection (0.072±0.032 h^-1^). In contrast, the lysogen displayed more moderate growth inhibition at both mid-infection (0.396±0.036 h^-1^) and late-infection (0.135±0.085 h^-1^), suggesting a degree of host immune protection. At the cellular level, principal component analysis of the proteomes showed a clear distinction based on strain type during mid- and late-, but not pre-infection, time points (**Fig. 1C**).

**Figure 1:**
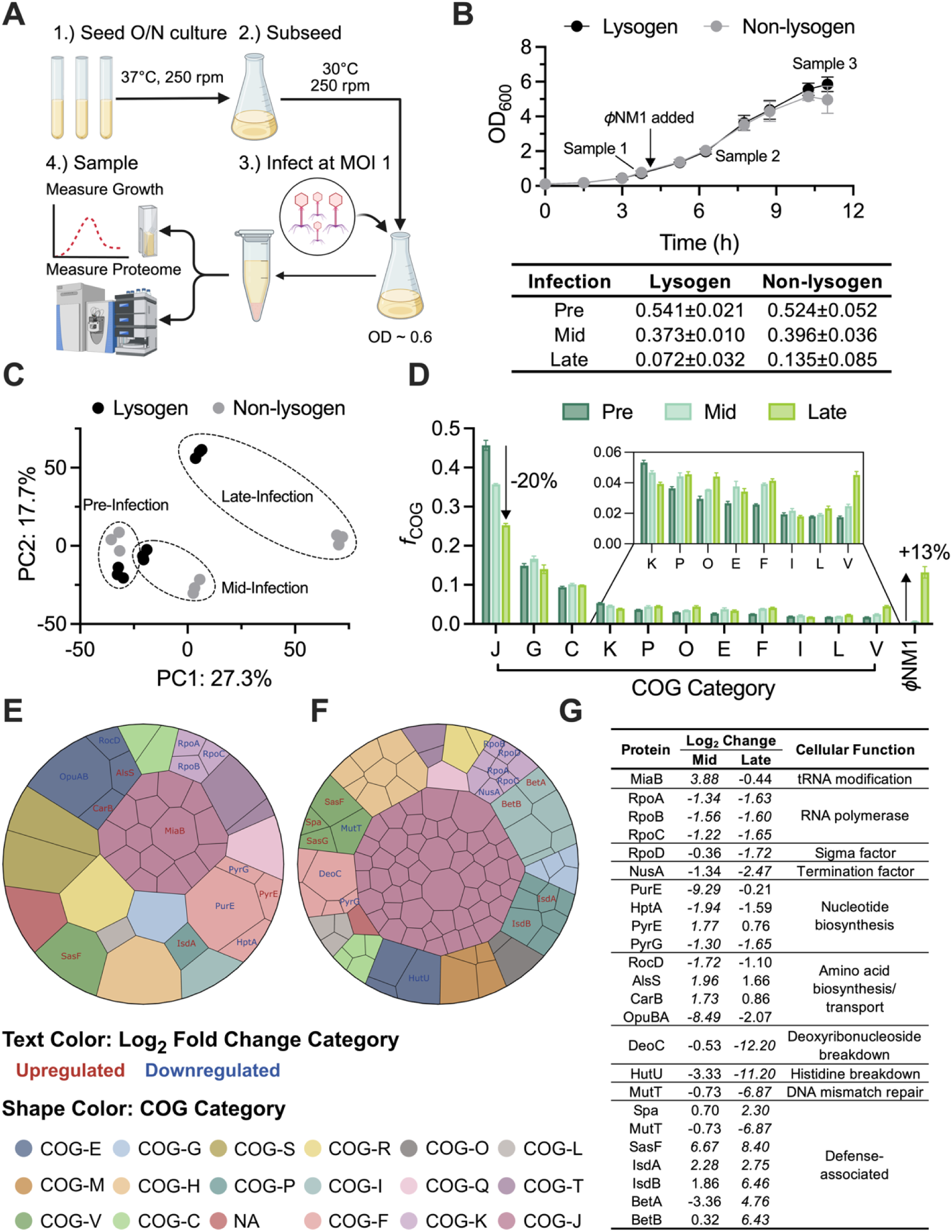
Lysogen and non-lysogen *S. aureus* strains exhibit divergent proteomes in response to phage infection. (**A**) Workflow for cell characterization, phage infection, and proteomics sample collection. (**B**) Growth curves and sampling time points for lysogen and non-lysogen cultures subjected to *ϕ*NM1 infection. Table shows mean values for maximum specific growth rates ± SD (n = 3). (**C**) Principal component analysis of proteomes from pre-, mid-, and late-infection time points. (**D**) Proteome fractions of top COG categories before and during *ϕ*NM1 infection. See **Dataset 1** for the complete list of COG categories. (**E–F**) Treemaps of DEPs during *ϕ*NM1 infection for (**E**) mid-infection vs. pre-infection and (**F**) late-infection vs. pre-infection comparisons. Blocks are colored by COG category and sized by absolute log_2_ fold changes. Text color indicates the category of log_2_ fold changes. **(G)** A representative list of DEPs in the non-lysogenic strain in response to *ϕ*NM1 infection. Italics numbers indicate a significant change.

### Longitudinal proteomes of the non-lysogen during ***ϕ***NM1 infection revealed significant resource reallocation resulting in growth inhibition

#### Phage infection caused translation shutdown of S. aureus, subsequent growth inhibition, and activation of host defenses

Proteome reallocation analyses(28, 29) were performed to assess which cellular processes were most affected in the non-lysogen by *ϕ*NM1 phage infection (**Fig. 1D, Dataset 1**). In the non-lysogen, pre-infection proteomes allocated 45.7% of all resources to Translation (COG-J), 14.9% to Carbohydrate Transport and Metabolism (COG-G), and 9.4% to Energy Generation (COG-C), with the remaining ∼30% spread across all other COGs (**Fig. 1D**). At mid-infection, investment into COG-J dropped by 10.0% from pre-infection levels while COG-G, Nucleotide Transport and Metabolism (COG-F), and Amino Acid Transport and Metabolism (COG-E) increased by 1.7%, 1.3%, and 1.1%, respectively (**Fig. 1D**). Increases in these key COGs could provide important resources for viral genome replication and protein production. All other COGs remained within 1% of their pre-infection values (**Fig. 1D**). At late-infection, investment into COG-J dropped by a further 10.4% for a total decrease of 20.4%, while COGs G, C, and E returned to within 1% of pre-infection values (**Fig. 1D**). COG-F increased by a further 0.3% for a cumulative increase of 1.6% (**Fig. 1D**). Besides COGs J and F, the only categories reallocated by more than 1% by the end of infection were Transcription (COG-K, - 1.4%), Posttranslational Modification, Protein Turnover, and Chaperones (COG-O, +1.4%), and Defense Mechanisms (COG-V, +2.8%) (**Fig. 1D**). Decreased abundance of two cold shock proteins (-0.88%) and RNA polymerase subunits (-0.82%) were the main drivers of COG-K suppression at late-infection, while COG-O mostly increased from chaperone expression: DnaK (+0.46%), ClpC (+0.28%), ClpP (+0.16%), GroEL (+0.15%). Higher investment in COG-V was primarily from increased expression of surface protein A (Spa, +1.33%) and alkyl hydroperoxide reductase subunit C (AhpC, +1.06%). In summary, *ϕ*NM1 infection resulted in large-scale shutdown of translation coupled with a smaller divestment from carbohydrate metabolism and transcription. Cells also made modest investments in nucleotide metabolism, chaperones, and defense mechanisms.

#### Analysis of differential protein expression across infection revealed the reprogramming of cellular processes in response to phage infection

Differentially expressed proteins (DEPs) were investigated during *ϕ*NM1 infection of the non-lysogen strain (**Fig. 1E, Dataset 2**). DEPs were classified as proteins with |log_2_ fold change| ≥ 1 and q-value *<* 0.05 relative to pre-infection levels. At mid-infection, there were a total of 41 host DEPs, with 13 being upregulated and 28 being downregulated (**Fig. 1E, Dataset 2**). Among these proteins, Translation (n = 15), Nucleotide Metabolism and Transport (n = 4), Amino Acid Metabolism and Transport (n = 4), and Transcription (n = 3) were the most enriched COGs (**Fig. 1E**). In Translation, 7/33 large subunits and 4/20 small subunits of the ribosome were downregulated, along with elongation factors Tu and G (**Fig. 1E**). The only upregulated COG-J protein was a tRNA methylthiotransferase, MiaB (**Figs. 1E, 1G**). 5-(carboxyamino)imidazole ribonucleotide mutase (PurE), hypoxanthine phosphoribosyltransferase (HptA), and CTP synthase (PyrG) were downregulated in COG-F while orotate phosphoribosyltransferase (PyrE) was upregulated (**Figs. 1E, 1G**). In COG-E, acetolactate synthase (AlsS) and carbamoyl-phosphate synthase large subunit (CarB) were upregulated and betaine/proline/choline family ABC transporter ATP-binding protein (OpuBA) and ornithine–oxo-acid transaminase (RocD) were downregulated (**Figs. 1E, 1G**). DEPs in COG-K included 3/6 subunits of the RNA polymerase (RpoA, RpoB, RpoC), which were all downregulated (**Figs. 1E, 1G**).

By late-infection, there were 113 host DEPs with 12 being upregulated and 101 being downregulated (**Fig. 1F, Dataset 2)**. Again, most DEPs were involved in Translation (n = 49), followed by Transcription (n = 8) and Coenzyme Transport and Metabolism (COG-H, n = 8) (**Fig. 1F**). Within Translation, 18/33 large subunits and 14/20 small subunits of the ribosome were downregulated, as were elongation factors Tu, G, and 4, initiation factor IF-3, peptide chain release factor 1, and alanine-, phenylalanine-, and leucine-tRNA ligases (**Fig. 1F**). There were no significantly upregulated proteins in Translation. In addition to RpoA, RpoB, and RpoC, transcription termination factor (NusA) and housekeeping sigma factor (RpoD) were also downregulated in Transcription (**Figs. 1F, 1G)**. Since RpoD is the primary sigma factor during exponential growth and directly regulates ribosomal proteins(30), its phage-responsive downregulation could help explain the drastic reduction in translation observed during infection (**Figs. 1D, 1E–1G**). The most downregulated proteins were deoxyribose-phosphate aldolase (DeoC, -12.2 log_2_) and urocanate hydratase (HutU, -11.2 log_2_), which are involved in the breakdown of deoxyribonucleosides and histidine, respectively (**Figs. 1F, 1G)**. Downregulation of DeoC and HutU could preserve important precursors for phage genome replication and protein production during the late stages of infection. Several defense-associated host genes were differentially regulated, including five upregulated surface proteins (Spa, SasF, SasG, IsdA, and IsdB) and two upregulated osmotic stress response genes (BetA and BetB, **Figs. 1F, 1G)**. Additionally, a homolog of the DNA mismatch repair protein MutT was downregulated (**Figs. 1F, 1G**).

Nearly all infection-related DEPs were downregulated, suggesting *ϕ*NM1’s primary infection strategy is to disrupt *S. aureus* host processes instead of co-opting them. This is consistent with previous reports demonstrating large-scale host transcript and/or protein knockdown in response to phage infection(31-33). Furthermore, inhibition of host translation has been reported for viruses of *Pseudomonas* and *Pseudoalteromonas* species, indicating that viruses may employ similar strategies for infecting target cells, even for highly divergent bacterial hosts(22, 34).

### Proteome dynamics revealed how *ϕ*NM1 hijacked host cellular processes and inactivated non-lysogenic *S. aureus* during infection

The *ϕ*NM1 proteome has 64 unique proteins, including 41 proteins with known functions, 10 proteins with putative functions, and 13 proteins with unknown functions (**Supplementary Table S1**). Before infection, no phage proteins were detected in the non-lysogen, as expected. At mid-infection, 30/64 phage proteins were detected and accounted for 0.7% of the whole proteome (**Figs. 2A, 2D, Dataset 1**). Of these proteins, the major head protein was by far the most abundant, accounting for 59.7% of all phage proteins (**Figs. 2A, 2C**). The major tail, antirepressor, phage portal, and upper and lower tail fiber proteins were next and contributed 11.7%, 6.4%, 4.2%, 2.9%, and 2.5% to the phage proteome, respectively (**Figs. 2A, 2C**). Next were several proteins that each contributed 1–2% to the phage proteome: a putative Arc-like repressor, head scaffolding protein, single-stranded DNA binding protein, dUTP diphosphatase, and two hypothetical proteins (**Figs. 2A, 2C**). At this point in *ϕ*NM1 takeover, endolysin was expressed at 0.4% of the phage proteome, but holin was not detected (**Fig. 2A**).

**Figure 2:**
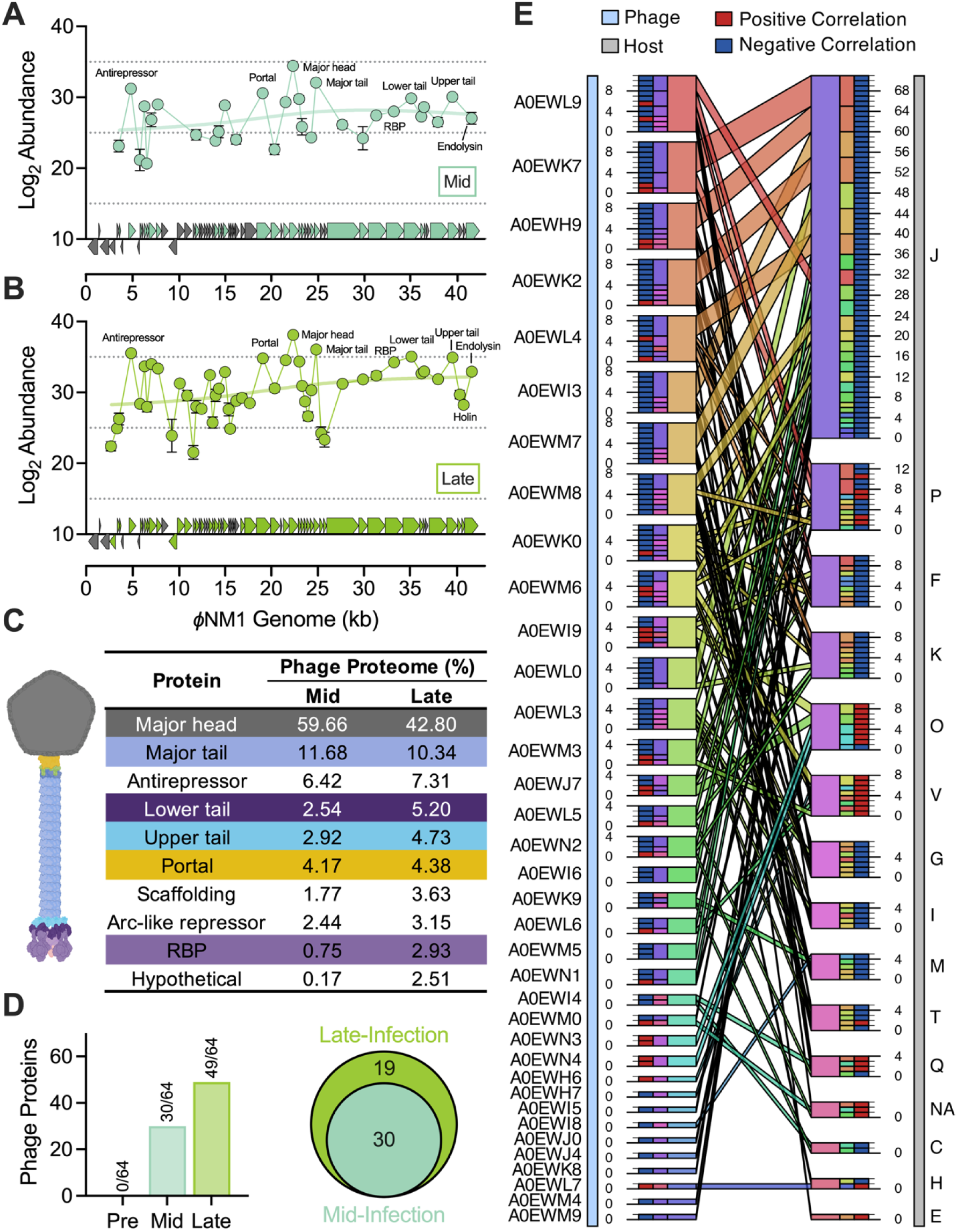
Proteomic profiles of *ϕ*NM1 and its dynamic interplay with non-lysogenic *S. aureus* during infection. (**A–B**) Protein abundances of non-lysogenic *S. aureus* at (**A**) mid- and (**B**) late-infection. (**C**) Number of detected *ϕ*NM1 proteins before and during infection and Venn diagram of *ϕ*NM1 proteins at mid- and late-infection. (**D**) Phage-host protein co-expression network for *ϕ*NM1 and non-lysogenic *S. aureus*. Each segment represents either a phage protein (left side) or a COG-based group of host proteins (right side), with segment size indicating the number of interactions. The innermost track provides a unique color for each segment, while the middle track summarizes interactions: phage protein segments show a functional breakdown, and COG segments detail interactions with phage proteins. The outermost track indicates whether the co-expression was positively (red) or negatively (blue) correlated. See **Dataset 1** for the complete list of COG categories.

Late in the infection cycle, 49/64 *ϕ*NM1 proteins were measured and comprised 13.2% of the whole proteome (**Figs. 1D, 2B, 2D, Dataset 1**). Again, the major head protein was the most abundant protein, representing 42.8% and 5.6% of the phage and total proteomes, respectively (**Figs. 2B, 2C**). Other structural proteins like the major tail, tail fibers, phage portal, and head scaffolding proteins were again highly expressed and together composed approximately 74% of the phage proteome. Interestingly, the 19 phage proteins expressed during late-, but not mid-infection, only accounted for 0.32% of phage proteins (**Figs. 2B–D**). DNA replication and packaging proteins were among the late-only proteins, including the DNA replication initiation proteins and the small and large terminase subunits. Surprisingly, the Cro and CI repressors were also found in this group. Although both Cro and CI were weakly expressed, Cro was 4.1-fold more abundant than CI, and with the coordinated action of the abundant antirepressor, was able to completely suppress integrase expression at both mid- and late-infection (**Figs. 2A, 2B**). Other late-only proteins included a Panton-Valentine leucocidin, two tail proteins (head-tail joining and tail completion), two tail assembly chaperones, and seven proteins of unknown or putative functions. Finally, holin was exclusively expressed at late-infection and accounted for 0.04% of the phage proteome, while endolysin increased from 0.4% to 1.2% of the phage proteome (**Figs. 2A, 2B**).

In summary, the dynamics of *ϕ*NM1’s proteome revealed the mechanisms of hijacking the host’s cellular processes for its reproduction cycle. During mid-infection, *ϕ*NM1 mainly used the commandeered resources to synthesize core structural components and the antirepressor. *ϕ*NM1 finished its infection cycle by replicating and packaging its DNA, completing construction of nascent virions, and lysing the cell through the activities of holin and endolysin. Overall, the phage primarily produced major structural proteins and the antirepressor during infection of the non-lysogen strain. DNA replication, packaging machinery, and holin were tightly regulated to only occur at the end of infection and in relatively low levels compared to structural proteins. This result supports a canonical holin-endolysin model of lysis wherein endolysin steadily accumulates in the cytoplasm throughout infection until holin reaches a critical pore-forming concentration, at which endolysin is released and cell lysis occurs(35).

#### Protein co-expression analysis revealed dynamic interactions between the host and ϕNM1

To investigate specific interactions between *ϕ*NM1 and *S. aureus*, a protein co-expression network was constructed for the most significant (p *<* 0.001) and highly correlated (|r| *>* 0.99) phage-host protein pairs (**Fig. 2E**). Even with such stringent criteria, there were a total of 154 co-expression relationships involving 36 phage proteins and 62 host proteins. COG-J showed the most interactions with phage proteins at 71, followed by Inorganic Ion Transport and Metabolism (COG-P) at 13, COG-F at 10, COGs K and O at 9, and COG-V at 8 (**Fig. 2E**). All host proteins in COGs J, F, and K were anticorrelated with their phage partners, but COG-O and COG-V host-phage relationships were mostly positive (**Fig. 2E**). Within COG-J, 77% of interactions were with ribosomal proteins and 11% were with elongation or initiation factors. Interactions with COGs K, O, and V were dominated by a select few proteins: 89% of COG-K interactions involved RNA polymerase subunit β’, 89% of COG-O interactions included Clp protease ClpA and foldase PrsA, and 75% of COG-V interactions were with surface protein SasG.

The phage protein A0EWL9 had the highest interaction count at 11, closely followed by A0EWK7 with 10, and then A0EWK2, A0EWL4, and A0EWH9, each with 9 interactions (**Fig. 2E**). A0EWL9 is the tail terminator protein, A0EWK7 is the transcriptional activator RinA, A0EWK2 is a dUTP diphosphatase, A0EWL4 is the major head protein, and A0EWH9 is the antirepressor. RinA is known to directly bind to the promoter region of host proteins and repress them^33^, so the observed anticorrelations with host proteins could represent direct interference. Most interactions with RinA were involved in Translation (n = 6), followed by Inorganic Ion Transport and Metabolism (n = 3), and Defense Mechanisms (n = 1) (**Fig. 2E**). Translation proteins were all anticorrelated and included two 50S subunits, three 30S subunits, and elongation factor G, while COG-P proteins were a putative phosphate transport regulator/metal ion binding protein (anticorrelated) and a GlyMet-binding protein (correlated). The sole COG-V protein was SasG and was positively correlated. SasG is a virulence-associated adhesin that has been shown to promote adherence to epithelial cells and biofilm formation(36).

Generally speaking, these data reveal that increasing levels of phage proteins correspond to decreasing levels of host housekeeping proteins, especially those in COGs J, F, K, and G (**Fig. 2E**). In contrast, host proteins known to be stress-inducible (e.g., those in COGs V and O) tend to increase as phage proteins increase (**Fig. 2E**). The expression of RinA is tightly coordinated with several host proteins, providing putative instances of direct, phage-mediated host repression that are hitherto unknown. Other phage proteins without DNA-binding domains may interact with host proteins directly (i.e., through protein-protein interactions) or correlate their expression through indirect mechanisms.

#### The economics of ϕNM1 infection in the non-lysogen highlight the host’s metabolic burden and growth inhibition

More than 20% reallocation of the proteome from Translation and other cellular processes toward 13% of the phage proteome suggests a significant energetic investment for phage generation (**Fig. 1D**). Accounting for the ATP costs of synthesizing amino acids, the reallocation from Translation freed 183.1±20.7 ATP equivalents that could then be used to fund phage production, which cost 176.3±21.7 ATP equivalents. These represent shifts of -16.1±1.6 and +12.7±1.5 percentage points, respectively, in the proteome-wide ATP budget from pre-infection levels. Assuming there are 1.35 x 10^6^ proteins per *S. aureus* cell (**Supplementary File S1**), translation shutdown freed 2.46 x 10^8^ ATP/cell and phage production demanded 2.37 x 10^8^ ATP/cell. In general, as infection proceeded, energetic demand increased, with pre-infection proteomes needing 1.72 x 10^9^ ATP/cell and late-infection proteomes needing 1.87 x 10^9^ ATP/cell, an increase of 1.5 x 10^8^ ATP/cell.

To investigate which amino reprogramming demanded the highest energetic cost, the amino acid fractions of the *ϕ*NM1 proteome were compared to those of the non-lysogen proteome at mid- and late-infection (**Fig. 3A–3C**). Proteomes of *ϕ*NM1 and non-lysogen required similar levels of amino acids during mid- and late-infection with R^2^ values of 0.81 and 0.83, respectively (**Fig. 3A, 3B**). Four amino acids were over 1% more utilized by *ϕ*NM1 at both mid- and late-infection, including asparagine, lysine, threonine, and phenylalanine (**Fig. 3A–3C**). Conversely, alanine, valine, and glycine were all over 1% or more utilized by the non-lysogen at mid- and late-infection (**Fig. 3A–3C**). Accounting for the ATP cost of each amino acid, lysine required the most ATP reallocation across infection, followed by threonine and asparagine (**Fig. 3D**). On average, *ϕ*NM1’s proteome was more energetically expensive than the host proteome at both mid-infection (+21.4±0.2 ATP equivalents) and late-infection (26.2±0.3 ATP equivalents) (**Fig. 3E**). Therefore, when switching from host to phage protein production, higher energy demands and different amino acid profiles necessitated reprogrammed metabolism and created metabolic burdens for the host, causing slowed growth (**Fig. 1B**).

**Figure 3:**
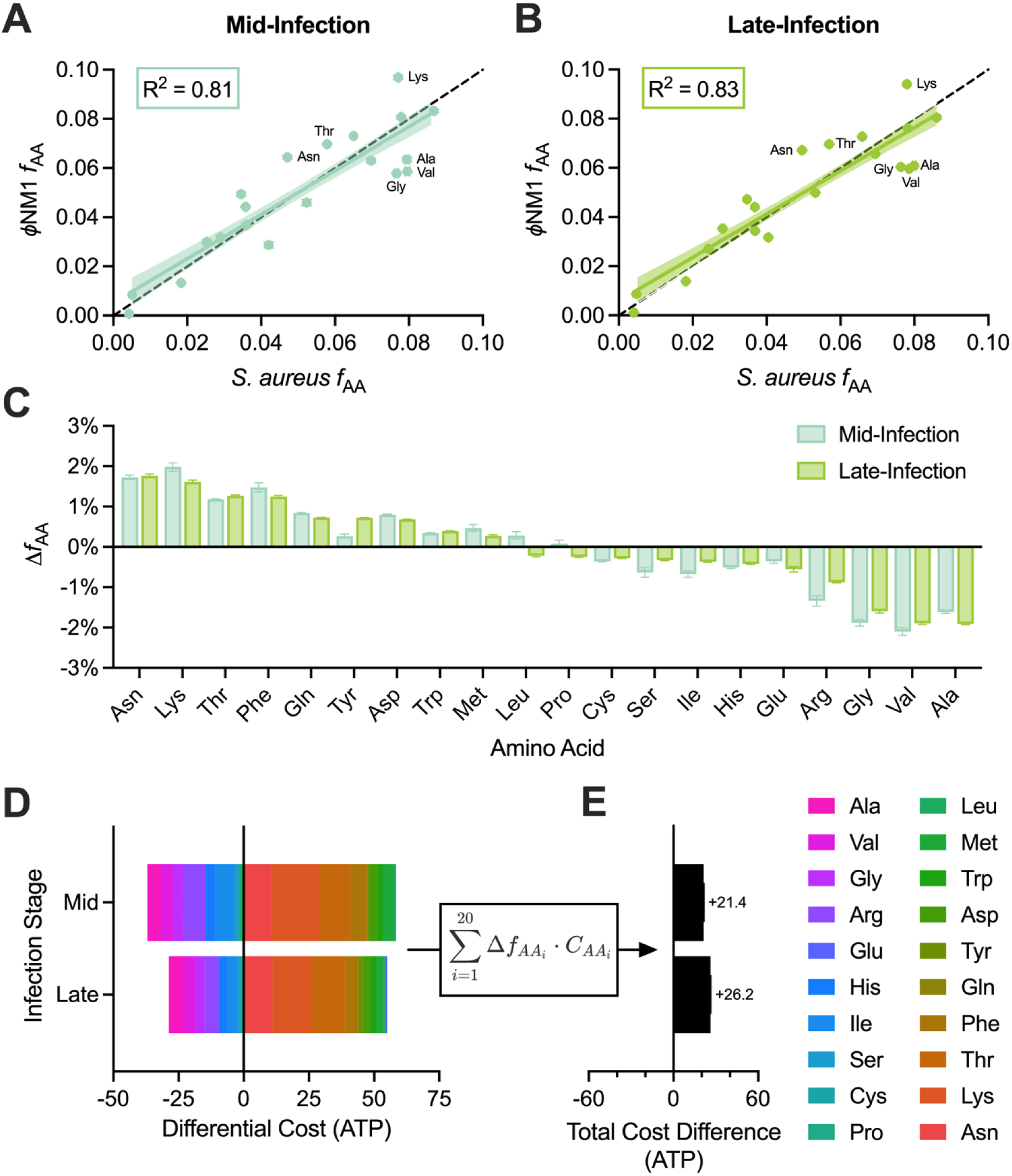
Economics of proteome reallocation during *ϕ*NM1 infection of non-lysogenic *S. aureus*. (**A–B**) Comparison of the amino acid fractions of *ϕ*NM1 and non-lysogen proteomes at (**A**) mid-infection and (**B**) late-infection. (**C**) Differences in the amino acid fractions of *ϕ*NM1 and non-lysogen during mid- and late-infection. (**D**) Differential cost of amino acids between *ϕ*NM1 and the non-lysogen per unit proteome. (**E**) Total cost difference between *ϕ*NM1 and non-lysogen per unit proteome during mid- and late-infection.

### Longitudinal proteomes elucidated the immunity of the lysogen to cope with *ϕ*NM1 infection

#### The lysogen boosted carbohydrate metabolism and defense systems while reducing translation shutdown and growth inhibition

We next sought to evaluate how a *ϕ*NM1 lysogen protected its host during superinfection by investigating longitudinal proteome allocation of the lysogen pre- and post-infection. During pre-infection, the lysogen strain apportioned proteomes like the non-lysogen (**Fig. 4A, Dataset 3**), and both exhibited a similar growth rate (**Fig. 1B**). Before infection, most protein investment of the lysogen was into COGs J, G, and C (44.3%, 14.8%, and 9.3%, respectively) (**Fig. 4A**), with a similar trend observed in the non-lysogen (**Fig. 1D**). COG-J decreased from pre- to mid-infection in the lysogen strain (**Fig. 4A**), but less than observed for the non-lysogen (-8.5% vs -10.0%) (**Fig. 1D, 4A**). From mid- to late-infection, COG-J dropped by a further 4.6% compared to 10.4% in the non-lysogen, resulting in a total decrease of 13.2% from pre-infection levels compared to 20.4% in the non-lysogen (**Fig. 1D, 4A**). COG-G increased from pre-infection to late-infection in the lysogen strain by 1.9% (**Fig. 4A**) compared to the small decrease observed for the non-lysogen (-0.9%) (**Fig. 1D**). Relative to pre-infection levels, the lysogen reallocated 7.3% and 2.8% more resources into COGs J and G, respectively (**Fig. 4A**), than the non-lysogen (**Fig. 1B**). This differential investment into COG-J was primarily due to increased production of elongation factors Tu and G (+1.97% and +0.74%, respectively), 30S ribosomal protein S5 (+0.51%), and 50S ribosomal protein L21 (+0.45%). Phosphopyruvate hydratase (+0.71%), fructose bisphosphate aldolase (+0.65%), transaldolase (+0.41%), and triose phosphate isomerase (+0.32%) were overproduced from COG-G.

**Figure 4:**
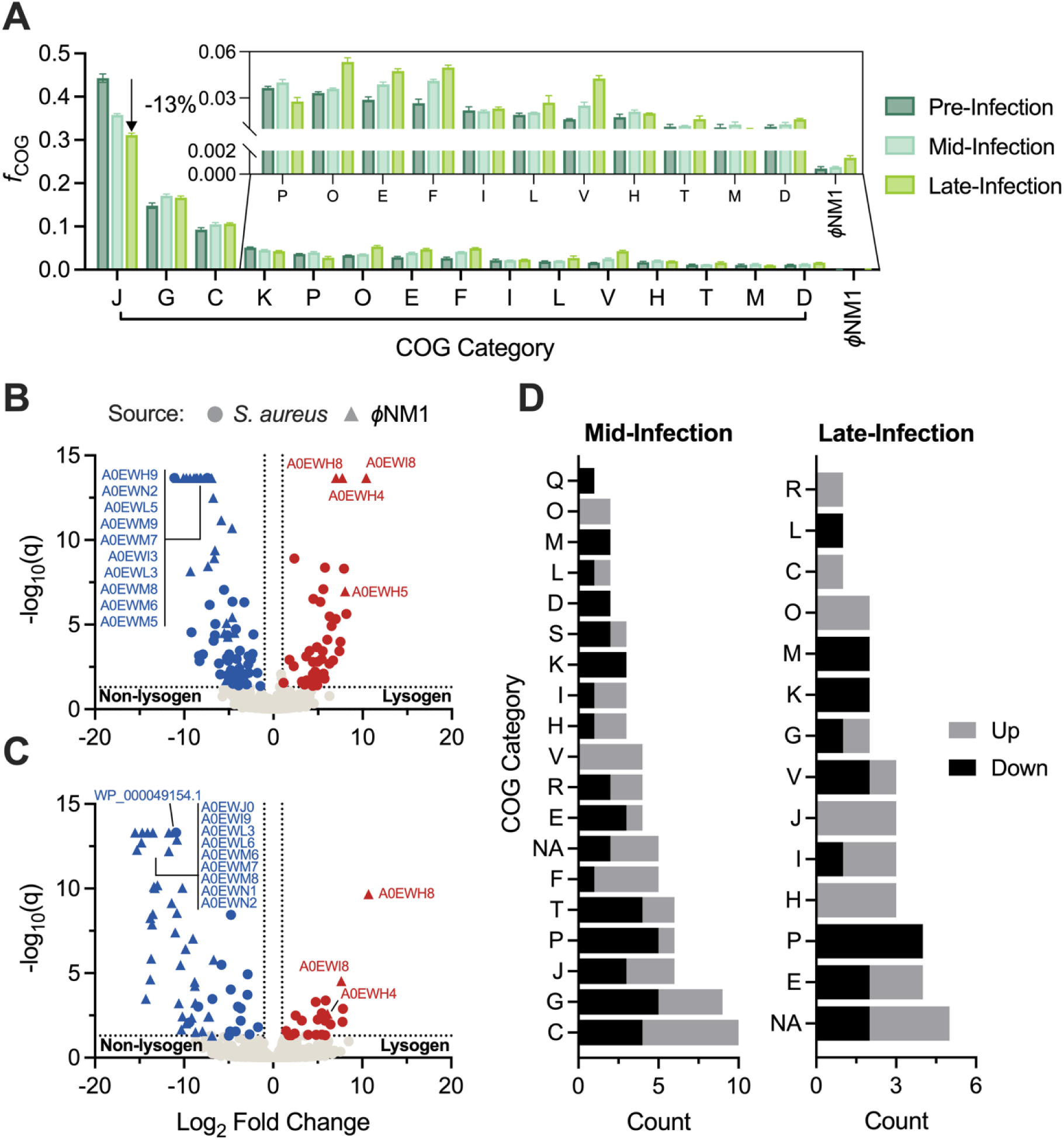
Proteome dynamics of lysogenic *S. aureus* in response to *ϕ*NM1 attack. (**A**) Proteome fractions of top 15 COG categories before and during *ϕ*NM1 infection. (**B–C**) Differential protein expression between lysogen and non-lysogen at (**B**) mid-infection and (**C**) late-infection. (**D**) Summary of DEPs between lysogen and non-lysogen by COG at mid- and late-infection. See **Dataset 1** for the complete list of COG categories.

Like the non-lysogen, the lysogen increased its proteome allocation in COG-V, gaining 0.9% by mid-infection and 2.7% by late-infection; however, Spa expression did not increase from pre-infection levels (**Fig. 4A**). Instead, resources were heavily funneled into AhpC (+0.75%) and protein deglycase (HchA, +0.33%). AhpC is an important player in oxidative stress defense(37), and HchA repairs proteins and nucleic acids from glycation arising due to reactive carbonyl species like methylglyoxal and glyoxal(38), whose production is encouraged by reactive oxygen species (ROS)(39). Notably, phage infection has been shown to generate ROS in a variety of bacteria, including *S. aureus*(40, 41), suggesting ROS defense may be critical for surviving viral attack.

Overall, these findings demonstrate that integrated *ϕ*NM1 successfully prevented host cell takeover by an invading phage, preserving resources in key cellular processes and shifting allocation of host defenses from virulence-associated surface proteins to ROS response.

#### Activation of stress responses and manipulation of amino acid pool helped the lysogen survive phage invasion

Differential expression analysis was performed to evaluate which host proteins were differentially regulated between the lysogen and non-lysogen in response to *ϕ*NM1 infection (**Figs. 4B–4D, Dataset 4**). Before infection, only two host proteins were significantly altered in expression between lysogen and non-lysogen, indicating the *ϕ*NM1 prophage did not heavily alter host protein expression under normal conditions (**Fig. 1C, Supplementary Fig. S1**). At mid-infection, 80 host proteins were significantly different between lysogen and non-lysogen, indicating divergent responses to *ϕ*NM1 attack (**Fig. 4B, 4D, Dataset 4**). DEPs were broadly distributed across COG categories, with 38 upregulated and 42 downregulated (**Fig. 4D, Dataset 4**). Several defense effectors (COG-V) were upregulated in the lysogen, including glutathione peroxidase (+3.24 log_2_ difference), a putative multidrug efflux ABC transporter (4.9 log_2_), restriction endonuclease subunit S (5.25 log_2_), and an ABC transporter ATP-binding protein (6.03 log_2_). This pattern of host defense activation is different from that of the non-lysogen, where surface proteins Spa and SasG dominated. Other stress-response associated DEPs included metalloprotease MroQ (+3.68 log_2_), clp protease subunit ClpL (+4.97 log_2_), and *clp* operon repressor CtsR (-7.41 log_2_). Collectively, these proteins influence quorum sensing and protein folding(42). Along with CtsR, two other transcriptional regulators were downregulated in the lysogen strain, both belonging to the MerR family, which are repressors known to respond to stress-related stimuli(43).

At late-infection, there were 36 host DEPs between the lysogen and non-lysogen; 19 were upregulated and 17 were downregulated in the lysogenic host (**Fig. 4C, 4D, Dataset 4**). Amino Acid Transport and Metabolism, Inorganic Ion Transport and Metabolism, and Coenzyme Transport and Metabolism had the most DEPs with four, four, and three, respectively (**Fig. 4D**). Focus was given to proteins involved in amino acid biosynthesis due to their expected importance in proteome reallocation. Among these were ornithine carbamoyltransferase (+7.83 log_2_ difference), urocanate hydratase (+7.80 log_2_), lysine-specific permease (-10.90 log_2_), and a His/Lys/Arg transporter (-4.76 log_2_). Decreased abundance of lysine permeases in the lysogen would result in less lysine, which was previously established to be critical for the *ϕ*NM1 proteome in the non-lysogen (**Fig. 3C**), while upregulation of ornithine carbamoyltransferase would increase arginine, which is the fourth most host-preferred amino acid (**Fig. 3C**). Pyridoxal 5’-phosphate synthase subunit PdxT was also upregulated in the lysogen (2.52 log_2_), which produces vitamin B6, a cofactor critical for the biosynthesis of various amino acids(44). Vitamin B6 has also been shown to quench ROS(45). Other proteins involved in oxidative stress response were upregulated, such as bacillithiol disulfide reductase (+5.9 log_2_) and thiol peroxidase (+1.45 log_2_).

Taken together, these findings suggest the lysogen mounts a more robust stress response to *ϕ*NM1 than the non-lysogen, especially toward ROS generated during infection(40, 41). Amino acid biosynthesis is directed away from amino acids most needed by *ϕ*NM1, like lysine, and is instead funneled into host-preferred amino acids like arginine (**Fig. 3C**). These metabolic adaptations prevent ribosome shutdown, as evidenced by the lack of downregulated Translation proteins at late-infection (**Fig. 4D**). Therefore, the lysogen specifically redirects host metabolism to repel *ϕ*NM1 invasion.

### Prophage-encoded defenses suppressed the expression of proteins from invading *ϕ*NM1

#### The ϕNM1 prophage expressed surveilling immune proteins before infection

Before infection, we detected 9/64 prophage proteins in the lysogen (**Fig. 5**). Almost all antisense genes were detected, including Tha-2 (A0EWH4), CI repressor (A0EWH5), A0EWH8, A0EWI0, and A0EWI8. Integrase (A0EWH2) was the only antisense-encoded protein not detected. The remaining four proteins were expressed from the positive strand and included the antirepressor, A0EWJ7, phage portal, and major head proteins. Like the closely related staphylococcal phage 80*α*, *ϕ*NM1 utilizes noncontiguous operons to facilitate lysis-lysogeny decisions, expressing primarily sense proteins during the lytic cycle and antisense proteins during the lysogenic cycle(46). Despite not being measured in our proteomics, integrase mRNA has been measured for 80*α* under lysogenic conditions(46), suggesting integrase repression during lysogeny may occur through an unknown post-transcriptional mechanism in these phages.

**Figure 5:**
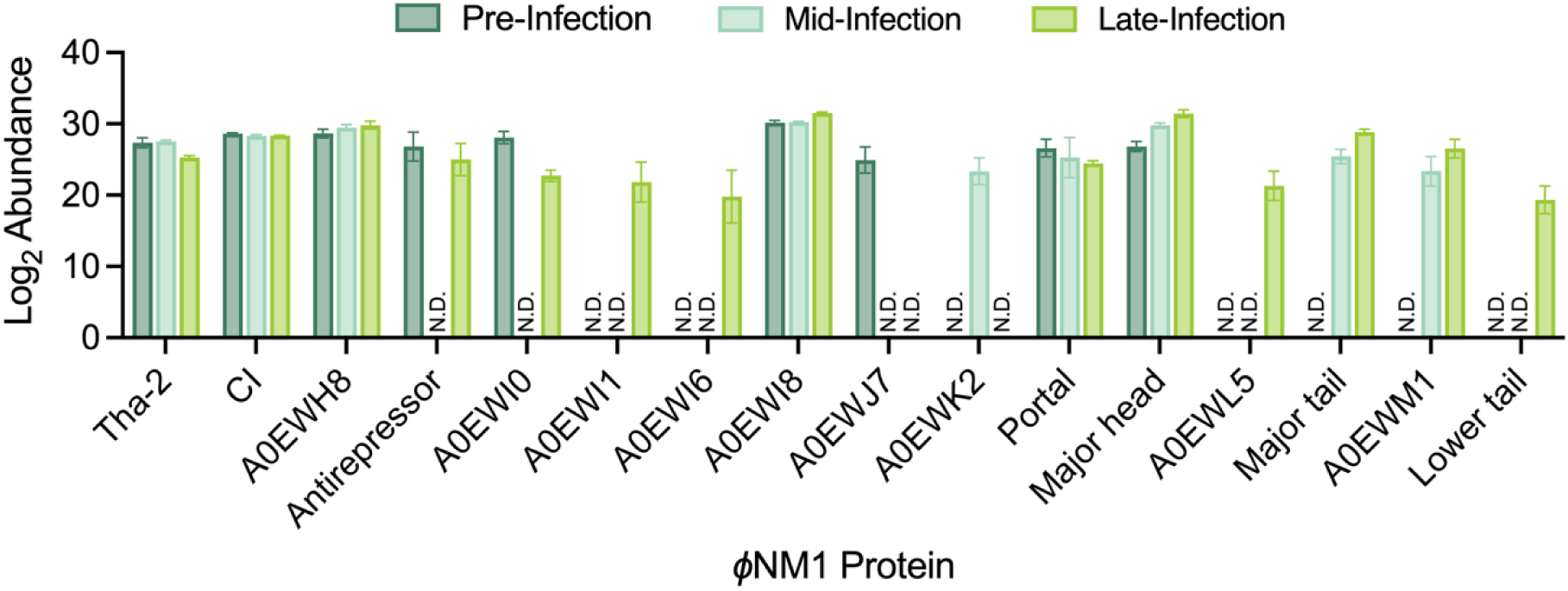
Phage protein expression before and during *ϕ*NM1 infection of lysogenic *S. aureus*.

The most plentiful protein at pre-infection was A0EWI8, which accounted for 42.2% of all phage proteins (**Fig. 5**). A0EWI8 is a DUF4393-containing protein, and DUF4393 originates from the Abi*α* protein of prophage 6 in *Enterococcus faecalis* V583(47). Currently, the mechanism of this protein remains unknown, but it is thought to result in premature lysis and non-productive infection of the invading phage. In *ϕ*NM1, A0EWI8 may work in concert with Tha-2 of the recently characterized Tha-ith abortive infection (Abi) system(46), which was also well-expressed (5.1% of prophage proteins). Tha-2 is a latent RNase that indiscriminately cleaves RNA and causes abortive infection when the lower tail fiber of invading phages is recognized. The second most abundant protein was A0EWH8, comprising 14.9% of the prophage proteome. It is an 81 amino acid uncharacterized protein that could not be annotated with either sequence- or structure-based homology searches and has no homolog in the well-studied staphylococcal phages 80*α*, *ϕ*11, or *ϕ*NM2. Along with Tha-2 and A0EWI8, A0EWH8 was among the most differentially upregulated phage proteins relative to the non-lysogen at both mid- and late-infection (**Fig. 5**). This finding, taken together with its similar genomic location and orientation to other phage immunity proteins, suggests A0EWH8 may play a role in blocking infection from competing phages. As expected, the CI repressor, which maintains lysogeny by preventing the transcription of lytic genes, was also strongly expressed under basal conditions (13.6% of prophage proteome).

Pre-infection expression of genes from the positive strand was mainly limited to proteins that were most prolific during productive *ϕ*NM1 infection of the non-lysogen, such as the major head, portal, and antirepressor proteins (**Fig. 5**). Given their high abundance during productive infection, expression from the prophage was likely due to leaky transcription, but could also signify secondary roles for these proteins during lysogeny. A0EWJ7, a Panton-Valentine leukocidin (PVL)-like protein, was also expressed despite being only moderately abundant during *ϕ*NM1 infection of the non-lysogen. Expression of the antirepressor was somewhat surprising given that some antirepressors directly inhibit the CI repressor^(48)^. In the more closely related staphylococcal phage *ϕ*11, however, the antirepressor enhances Cro repressor binding to outcompete the CI repressor(49). Since antirepressor activity depends on the presence of Cro and no Cro protein was detected pre-infection, low levels of antirepressor expression are likely tolerated under lysogeny conditions by *ϕ*NM1.

At mid-infection, 9/64 *ϕ*NM1 proteins were measured, and only three *ϕ*NM1 proteins significantly changed in expression. The major head protein was upregulated (2.97 log_2_), presumably from infecting phage, and the antirepressor and A0EWI0 (function unknown) were downregulated (-6.42 and -8.32 log_2_, respectively). Both the antirepressor and A0EWI0 were not detected in mid-infection samples. Additionally, A0EWJ7 was not measured at mid-infection. Immunity-associated proteins Tha-2 and A0EWI8 did not significantly change in abundance, suggesting basal expression is sufficient for mediating immunity. At this point, A0EWM7, the lower tail fiber protein that activates Tha-2-mediated immunity, was not detected, nor was the Tha-2 inhibitor, Ith(46). Three phage proteins not measured pre-infection were detected at mid-infection: dUTP diphosphatase, major tail protein, and tail chaperone protein (A0EWM1). A0EWI8 remained the most abundant phage protein (34.06% of phage proteome), followed by the major head protein (25.46%), A0EWH8 (20.92%), and the CI repressor (9.30%). In total, *ϕ*NM1 proteins accounted for 0.06% of the measured proteome at this stage of infection, compared to 0.66% when infecting the non-lysogen strain, demonstrating that the *ϕ*NM1 prophage can suppress invading phage proteins by over 10-fold in the middle stages of infection.

At late-infection, 14/64 *ϕ*NM1 proteins were expressed and none significantly changed in expression from mid-infection. Once again, A0EWI8 and the major head protein were most highly expressed (38.08% and 35.52% of the phage proteome, respectively). A0EWH8, the major tail protein, and the CI repressor rounded out the top five most abundant *ϕ*NM1 proteins at 11.87%, 6.61%, and 4.49% of the phage proteome, respectively. A0EWI0 was the only protein expressed at mid-infection that was not also expressed at late-infection, and except for the antirepressor, the proteins expressed at late-, but not mid-infection, were of extremely low abundance. In total, *ϕ*NM1 proteins accounted for 0.14% of the measured proteome at late-infection, compared to 13.2% when infecting the non-lysogen strain. In summary, invading *ϕ*NM1 was unable to hijack host resources and create virions in the lysogen strain due to defensive prophage proteins.

### Coinfection with phage-like particles and *ϕ*NM1 induced unique metabolic reallocation and accelerated the phage lytic cycle in non-lysogenic *S. aureus*

CRISPR-Cas antimicrobials offer precision therapies for treating *S. aureus*(14, 18, 19). These antimicrobials are phage-like particles containing CRISPR-Cas systems that recognize, target, and disrupt a gene of interest. This genetic interference results in chromosomal injury and inactivation of gene functions, ostensibly causing cell death. These treatments are generally produced through the induction of a lysogenic strain carrying the phage-based antimicrobial. Due to the presence of the packaging signal on the resident prophage, production of CRISPR-Cas antimicrobials typically includes a mixture of wildtype phage and CRISPR-Cas systems. It is not well-understood how the introduction of orthogonal DNA and the associated expression of proteins for replication, selection, and interference impact the phage infection process. To address these knowledge gaps, we engineered CRISPR-carrying phage-like particles and evaluated their effect on the infection of lysogenic and non-lysogenic *S. aureus*.

#### ϕDD0001 was more lethal to the non-lysogen than ϕNM1, but did not significantly increase metabolic burden

A null-targeting CRISPR plasmid was constructed by placing *Streptococcus pyogenes* Cas9 (SpCas9) under the constitutive P*_rpsL_* promoter and an empty guide RNA cassette under the P*_cap1A_* promoter. For phagemid packaging, *terS* from *ϕ*NM1 was cloned into the CRISPR plasmid without a promoter. A null-targeting system was selected to prevent artifacts arising from chromosomal injury and disruption of gene function, instead focusing on the metabolic burden associated with expressing phagemid-encoded proteins. In total, three proteins were expected to be expressed in *S. aureus*: SpCas9, the *S. aureus* plasmid replication protein (RepF), and a chloramphenicol resistance marker (CAT). A mixture of phage-like particles and wildtype phage (*ϕ*DD0001) was generated by infecting the lysogen harboring the phagemid with purified *ϕ*NM1. Both the lysogenic and non-lysogenic *S. aureus* strains were infected with *ϕ*DD0001, and cell growth kinetics and proteomes were analyzed (**Fig. 6**).

**Figure 6:**
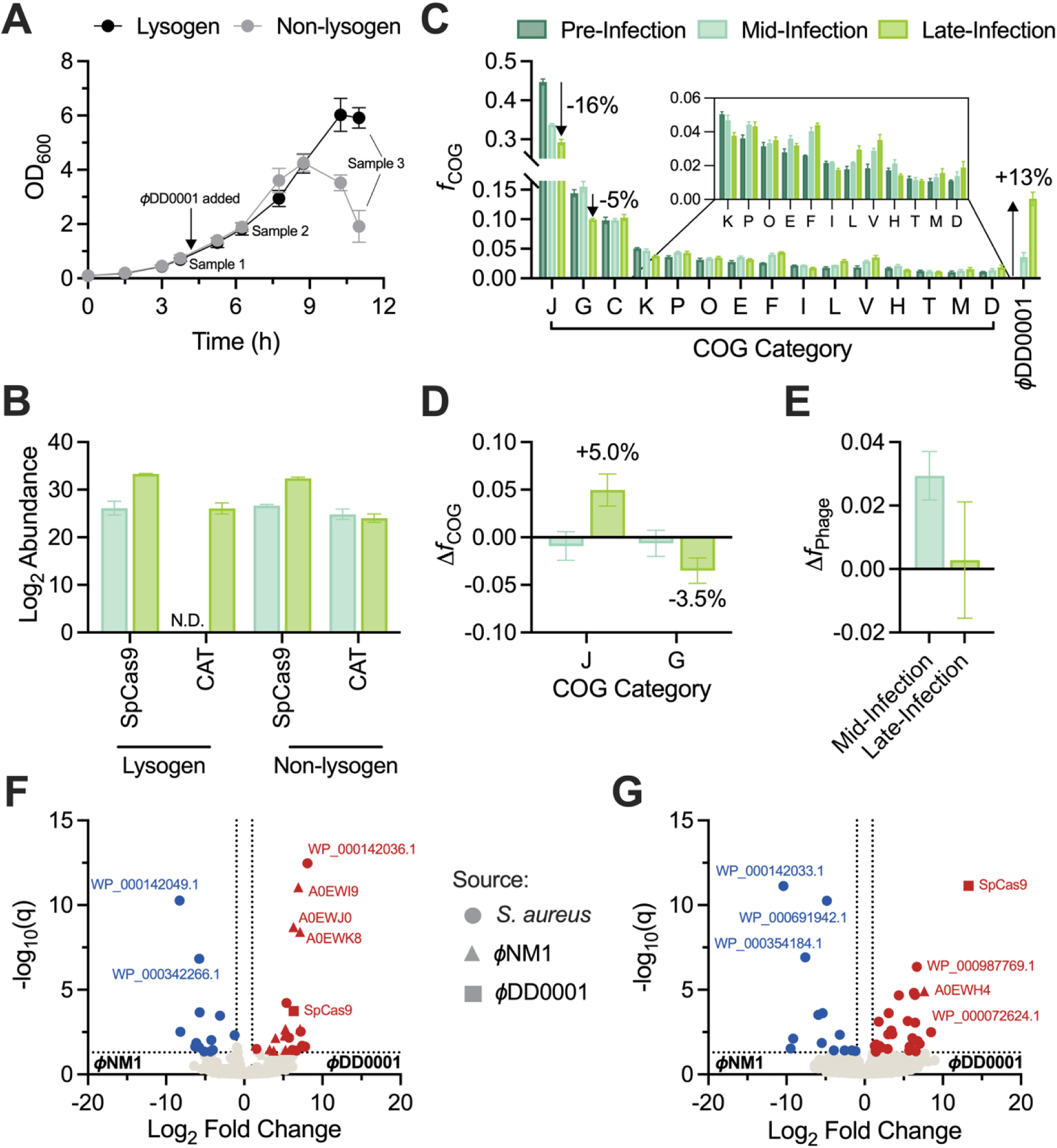
Engineered phage-like particles modulate host and phage proteomes differently than wildtype *ϕ*NM1 during non-lysogen infection. (**A**) Growth curves and sampling time points for lysogen and non-lysogen cultures subjected to *ϕ*DD0001 infection. (**B**) Expression of phagemid-encoded proteins during *ϕ*DD0001 infection of lysogen and non-lysogen. (**C**) Proteome fractions of top 15 COG categories before and during *ϕ*DD0001 infection of non-lysogen. (**D**) Change in proteome reallocation for COGs J and G during infection of non-lysogen with *ϕ*DD0001 vs. *ϕ*NM1. (**E**) Change in proteome fractions of *ϕ*DD0001 vs. *ϕ*NM1 during non-lysogen infection. (**F–G**) Differential protein expression between *ϕ*DD0001- and *ϕ*NM1-infected non-lysogenic *S. aureus* at (**F**) mid-infection and (**G**) late-infection. See **Dataset 1** for the complete list of COG categories.

In the non-lysogen, *ϕ*DD0001 initiated culture collapse two hours before *ϕ*NM1, despite infecting at the same MOI and cell density (**Figs. 1A, 6A**). Yet, mid-infection growth rates between *ϕ*DD0001- and *ϕ*NM1-infected cells were remarkably similar (**Supplementary Fig. S2A**). By late-infection, however, *ϕ*DD0001 achieved an apparent killing rate of -0.333±0.148 h^-1^, which was significantly lower (more lethal) than *ϕ*NM1 (**Supplementary Fig. S2A**). Growth rates of lysogenic *S. aureus* did not change based on phage treatment for any infection stage, indicating *ϕ*DD0001 did not cause a significant metabolic burden and prophage defenses remained active against contaminating *ϕ*NM1 particles (**Supplementary Fig. S2B**).

#### Phage-like particles delivered cargos, evaded prophage and host defenses, and expressed therapeutic proteins

Given the strong repression of most phage proteins during *ϕ*NM1 infection of the lysogen (**Fig. 4**), we first asked whether phagemids delivered by *ϕ*DD0001 were able to evade prophage defenses in the lysogen and express their proteins. At mid-infection, we observed expression for SpCas9 (26.1 log_2_ abundance), but not CAT or RepF (**Fig. 6B**). By late-infection, CAT levels were detectable (26.1 log_2_) and SpCas9 had increased by 7.2 log_2_ to 33.3 log_2_. Again, no RepF was detected. Only 7/64 and 14/64 *ϕ*NM1 proteins were detected at mid- and late-infection, respectively, indicating the lysogen consistently suppressed phage proteins whether infected with *ϕ*NM1 or *ϕ*DD0001. Therefore, *ϕ*DD0001 was able to successfully deliver, evade, and express its cargo despite active prophage defenses. Tha-2 is expected to be the main prophage defense and is activated by A0EWM7(46). The expression of phagemid, but not phage, proteins suggests phage genomes are not co-packaged in the same capsid with phagemids, as injection of both would result in A0EWM7 production, Tha-2 activation, and non-specific degradation of phagemid and phage mRNAs. In this model, *ϕ*DD0001 contains a mixture of pure wildtype phage particles and pure engineered phage-like particles as opposed to particles with chimeric or co-packaged DNAs.

During productive infection of the non-lysogen, phagemid proteins were again expressed (**Fig. 6B**). SpCas9 increased from 26.6 to 32.4 log_2_ across infection for a total change of +5.8 log_2_, CAT remained relatively constant at 24.8 and 24.0 log_2_, and RepF was again undetected (**Fig. 6B**). Phagemid protein production did not significantly vary by strain, underscoring the minimal interference of prophage defenses on phagemid protein expression (**Fig. 6A, 6B**).

In summary, a mixture of wildtype and engineered phage-like particles was able to successfully deliver and express nucleic acid cargos in both lysogenic and non-lysogenic *S. aureus*. The robust and persistent expression of SpCas9 in the lysogen strain is especially notable because it demonstrates the ability of CRISPR-carrying phage-like particles to evade host and prophage immune systems proven to limit expression of normal phage DNA. Next, we elucidated the features of *ϕ*DD0001 that led to improved killing kinetics in the non-lysogen using proteome reallocation and differential expression analyses.

#### ϕDD0001 prioritized the shutdown of glycolysis over translation and prevented host stress response activation

Consistent with the observed growth rates, at mid-infection, non-lysogen proteomes infected with *ϕ*DD0001 were similar to those infected with *ϕ*NM1 (**Figs. 1D, 6C, Dataset 5**). COGs J and G were reduced by 1.9% and 1.1% relative to *ϕ*NM1, respectively, but no other COGs exceeded a differential allocation of 1% (**Fig. 6C, 6D**). By late-infection, the host proteome had undergone significant reallocation. *ϕ*DD0001-treated cells reallocated COG-J by -15.5% and COG-G by -4.5%, which resulted in 5.0% more COG-J and 3.5% less COG-G than *ϕ*NM1 (**Fig. 6C, 6D**). The relative increase in COG-J was almost entirely due to elongation factor Tu (+3.80%), with modest contributions from 50S ribosomal proteins L5 and L21 and translation initiation factor IF-2 (+0.27%, +0.24%, and +0.19%, respectively). The relative decrease in COG-G primarily came from fructose bisphosphate aldolase (-1.33%), type I glyceraldehyde-3-phosphate dehydrogenase (-0.78%), and phosphopyruvate hydratase (-0.36%), which are major glycolytic enzymes. Proteome reallocation into stress-implicated COGs V and O was 1.08% and 1.03% less for *ϕ*DD0001 than for *ϕ*NM1. In COG-V, AhpC and Spa were the most differentially reallocated (-0.55% and -0.48%, respectively), while in COG-O it was chaperones DnaK, GroEL, and ClpP (-0.26%, -0.15%, and -0.11%, respectively), as well as trigger factor (-0.15%), accounting for most of the difference. Next, DEPs between phage treatments at mid- and late-infection were evaluated.

There were 25 host DEPs between *ϕ*DD0001 and *ϕ*NM1 during mid-infection, with 12 upregulated and 13 downregulated (**Fig. 6F**, **Dataset 6**). Notable DEPs included two stress response regulators, YycF (+6.69 log_2_ difference) and CtsR (-5.71 log_2_), and two proteins involved in DNA Replication, Recombination, and Repair (COG-L): DNA primase (+7.78 log_2_), and ribonuclease HII (-5.13 log_2_). At this stage, there were no DEPs from COG-G and only four DEPs from COG-J, consistent with proteome reallocation analyses. During late-infection, there were 41 host DEPs, with 27 upregulated and 14 downregulated (**Fig. 6G**, **Dataset 6**). Consistent with proteome reallocation analysis (**Fig. 6C, 6D**), COG-J was the most affected COG category with seven DEPs, all of which were upregulated in *ϕ*DD0001. COG-L was the second most affected COG with four DEPs: ribonuclease HII (+8.49 log_2_), segregation/condensation protein A (+6.55 log_2_), DNA polymerase III subunit ⍺ (+2.94 log_2_), and DNA helicase PcrA (+1.90 log_2_). One stress-related repressor was significantly downregulated (WP_000160435.1, -5.49 log_2_), as were two paralogs that repair oxidative damage to proteins (WP_000159902.1, -2.57 log_2_ and WP_001024830.1, -2.44 log_2_). No glycolysis proteins were among the DEPs, implying the -3.5% change in COG-G was due to the aggregation of many small-scale protein downregulations.

Overall, *ϕ*DD0001 was able to achieve greater killing rates than *ϕ*NM1 in non-lysogenic *S. aureus* by prioritizing shutdown of glycolysis over translation and mitigating reallocation into host defenses. Compared to *ϕ*NM1, *ϕ*DD0001 also downregulated key stress effectors and upregulated DNA replication machinery, which likely accelerated phage genome replication, protein production, and lytic activity.

#### Earlier phage replication led to accelerated cell death in non-lysogenic S. aureus

Unlike the host proteome, significant differences in phage proteome expression were observable by mid-infection. *ϕ*DD0001 accounted for 3.6% of the total proteome at mid-infection compared to 0.7% for *ϕ*NM1, a change of +2.9% (**Fig. 6E**). The main proteins with increased proteome fraction were the major head (+1.5%), major tail (+0.25%), tail fiber (+0.17%), and phage portal (+0.17%) proteins. At late-infection, both phages occupied ∼13% of the measured proteome and differed by only 0.3%, indicating that the total amount of phage proteins produced during infection did not significantly change (**Figs. 1D, 6C**). Phagemid-encoded proteins did not significantly contribute to the protein fractions, at most accounting for 0.09% of the proteome.

There were 12 mid-infection phage DEPs, and all were upregulated in *ϕ*DD0001 relative to *ϕ*NM1 (**Fig. 6F, Dataset 6**). Most of the proteins were tail structural proteins, including the head-tail connector (A0EWL6, +4.01 log_2_), head-tail joining (A0EWL7, +5.33 log_2_), tail completion (A0EWL8, +5.28 log_2_), tail terminator (A0EWL9, +3.75 log_2_), distal tail (A0EWM4, +5.22 log_2_), and tail-associated lysin proteins (A0EWM5, +3.25 log_2_). Also upregulated were two hypothetical proteins, the ejection protein (A0EWL1, +5.28 log_2_), small terminase (A0EWK8, +7.13 log_2_), A0EWI9 (+6.93 log_2_), and A0EWJ0 (+6.31 log_2_). The small terminase was used as a packaging signal for the CRISPR-Cas phagemid, which likely contributed to its high expression, although it was not placed under a promoter. A0EWI9 and A0EWJ0 are homologs of ORFs 20 and 21 in staphylococcal phage 80*α*, respectively, and together initiate phage DNA replication, likely through recruitment of the host DNA helicase(50). For *ϕ*NM1, A0EWI9 and A0EWJ0 were only detected during late-infection, which together with increased tail protein expression, implies *ϕ*DD0001 progressed through the lytic cycle more rapidly than *ϕ*NM1. Upregulation of phage DNA replication likely created more templates for mRNA synthesis, accelerating the assembly of phage particles. This hypothesis is supported by the upregulation of members of the host replisome at different stages of infection, including DNA primase, DNA helicase, and DNA polymerase III subunit ⍺. At late-infection, the only phage protein differentially expressed was Tha-2 (+7.64 log_2_) (**Fig. 6G, Dataset 6**).

In summary, host resources were more quickly reallocated toward phage proteins in *ϕ*DD0001-infected cells, but final resource investment into phage proteins was similar for both phage treatments. Accelerated production of phage proteins coincided with earlier expression of phage DNA replicative machinery and overexpression of the host replisome. Together, these data help explain how an engineered phage variant was able to achieve more favorable infection kinetics than its natural counterpart and provide a framework for understanding and optimizing engineered phage therapies.

## Conclusion

This study provides a systems-level understanding of the intricate interplay between phage *ϕ*NM1 and its *S. aureus* host, which is crucial for designing next-generation precision phage therapies. Proteome reallocation elucidates the dynamics and mechanisms of phage infection and defense in *S. aureus*. Phage infection caused more severe growth defects in non-lysogenic *S. aureus* than in the lysogenic strain. Both the lysogen and non-lysogen exhibited distinct proteomic profiles during phage infection, with up to 20% reduction in proteome allocation toward protein translation in the non-lysogen. During infection of the non-lysogen, *ϕ*NM1 expressed approximately 77% of its proteome, hijacking over 13% of the host’s proteome investment and activating phage replication and lysis. The economics of *ϕ*NM1 in the non-lysogen highlights the host metabolic burden and growth inhibition. However, during infection of the lysogen, *ϕ*NM1 expressed only about 22% of its proteome, hijacking just 0.14% of the host’s proteome investment, and failing to activate phage replication and lysis. Overexpression of host and phage defense proteins, along with reprogrammed metabolism in the lysogen, provided immunity to phage infection and prevented growth collapse. Furthermore, coinfection with engineered phage-like particles and wildtype phage triggered a unique proteomic and metabolic reallocation, resulting in enhanced cell killing kinetics. We anticipate the findings will aid the rational design of phage therapies, in which phages can specifically recognize their host, inject cargos with desirable therapeutics, evade host defenses, and activate the cargoes for targeted therapeutic treatment against multi-drug resistant pathogens. Such cargoes, including CRISPR-Cas antimicrobials with high specificity, efficiency, adaptability, and modularity in design(51), can provide powerful toolboxes for developing next-generation precision therapeutics

## Materials and Methods

### Strains, plasmids, phagemids, and phages

#### Strains

All strains used in this study are found in **Supplementary Table S2**. *S. aureus* RN4220 (#NR-45946) was obtained from BEI, and *S. aureus* Newman was obtained from ATCC (#25904). To construct the lysogen strain SaDD0001, purified *ϕ*NM1 was used to infect *S. aureus* RN4220, and turbid plaques were restreaked on tryptic soy agar (TSA) and grown overnight at 37℃. PCR was used to confirm the presence of the integrated prophage, yielding strain SaDD0001.

#### Plasmids and phagemids

All plasmids and phagemids used in this study are found in **Supplementary Table S2**. The pCasSA editing plasmid (Addgene #98211) was a gift from Quanjiang Ji and served as the backbone for the empty phagemid pDD0001(52). pDD0001 was constructed using Gibson assembly, as previously described(53). Briefly, pCasSA was amplified with pCasSA.f/r and the *ϕ*NM1 *terS* gene, which contains the *pac* site, was amplified using Phusion™ Hot Start II DNA Polymerase (Thermo Scientific™ #F549L) according to the manufacturer’s recommendations. Assembly reaction mixtures were heat-shock transformed into *E. coli* DH10β and clones were confirmed via colony PCR and Sanger sequencing. pDD0001 was transformed into *S. aureus*, as previously described(54) and confirmed via colony PCR.

#### Phages

All phages used in this study are found in **Supplementary Table S2**.

### Cell culture

#### Media

*E. coli* strains were grown in lysogeny broth (LB, including 10.0 g/L tryptone, 5.0 g/L yeast extract, 10.0 g/L NaCl) with 50 *µ*g/mL kanamycin when appropriate. *S. aureus* strains were grown in tryptic soy broth (TSB, including 17.0 g/L tryptone, 5.0 g/L NaCl, 3.0 g/L soy peptone, 2.5 g/L glucose, and 2.5 g/L dipotassium phosphate) with 10 *µ*g/mL chloramphenicol when appropriate. During phage infection, cultures were supplemented with 5 mM CaCl_2_.

#### Culturing conditions

Unless otherwise noted, all strains were grown at 37 °C with 250 rpm shaking in a Thermo Scientific MaxQ 6000 incubator.

### Phage methods

#### Isolation of ϕNM1 from S. aureus Newman

To begin, an overnight culture of *S. aureus* Newman was diluted 50x into fresh media and grown to mid-exponential culture before induction with 2 µg/mL mitomycin C. Induced cultures were incubated for 6 hr, followed by the addition of 2 drops of chloroform to kill surviving cells. Cultures were incubated for 15 min at 37 ℃ with shaking and then spun down to pellet cell debris. Supernatant was decanted into a new tube with care not to disturb the cell debris, and 1 additional drop of chloroform was added. Supernatants were filtered through a sterile 0.22 µm membrane. Filtered Newman lysate was used to infect *S. aureus* RN4220, then isolated plaques were picked, phage particles were freed from the agar, and phage-specific primers were used to PCR the resuspended phage to determine which phages were present (**Supplementary Table S2**). Samples that were PCR positive for *ϕ*NM1 were then used to re-infect *S. aureus* RN4220, and the process was repeated a total of three times. The final purified phage was PCR positive for *ϕ*NM1, but not *ϕ*NM2–4.

#### Lysogenization of S. aureus RN4220

Purified and isolated *ϕ*NM1 was spotted on a lawn of *S. aureus* RN4220, and plaques were allowed to develop overnight at 37 ℃. Spots with turbid centers were restreaked on fresh TSA plates and grown overnight at 37 ℃. Colony PCR with phage-specific primers was used to determine which, if any, prophages had integrated into the strain (**Supplementary Table S2**). *S. aureus*-specific primers were included as a positive control. Colonies PCR positive for *ϕ*NM1, but not *ϕ*NM2–4, were saved and confirmed for lysogeny by resistance to superinfection by *ϕ*NM1 (no plaque phenotype).

#### Generation of purified phage stocks

Purified *ϕ*NM1 and *ϕ*DD0001 were generated by growing *S. aureus* RN4220 either with the pDD0001 phagemid (*ϕ*DD0001) or without (*ϕ*NM1) to mid-exponential phase before infecting with previously isolated *ϕ*NM1 at MOI ∼ 1. *S. aureus* RN4220 harboring pDD0001 was grown under selection at 30 ℃ to avoid plasmid loss(52). Once infected, cultures were incubated overnight at 30 ℃, 80 rpm to allow time for lysis. Cleared cultures were centrifuged, and the supernatant was collected and filtered through a sterile 0.22 µm membrane.

#### Determination of plaque titers

To titer phage lysates, a lawn of *S. aureus* RN4220 was first prepared by mixing 100 µL of overnight culture with 4.9 mL of 0.7% soft tryptic soy agar (TSA) supplemented with 5 mM CaCl_2,_ pouring the culture-agar mixture over a 1.5% TSA plate, and allowing the lawn to solidify for ∼5 min. Filtered lysates were serially diluted, and 3 µL was spotted on the *S. aureus* lawn. Spots were allowed to dry for 5–10 min before plates were placed lid side up for overnight incubation at 37 ℃. The following day, plaques were enumerated and used to calculate PFU/mL.

#### Phage infection experiments

Both the non-lysogenic strain RN4220 and lysogenic strain RN4220*^ϕ^*^NM1^ (or SaDD0001) were grown in shake flasks and infected at an MOI of 1 based on PFU with either pure *ϕ*NM1 or *ϕ*DD0001—a mixture of *ϕ*NM1 and *ϕ*NM1-based phage-like particles—at exponential growth phase (OD_600nm_ = 0.6–0.9) (**Figs. 1A, 1B**). We performed experiments for each culture (i.e., RN4220 or SaDD0001) and each treatment (i.e., *ϕ*NM1 or *ϕ*DD0001) in biological triplicates and collected samples for LC-MS/MS-based proteomics at three timepoints: pre-infection (t = 0 hr), mid-infection (t = 2.5 hr), and post-infection (t = 7.5 hr). Cells were spun down, washed once with sterile Millipore water, and pelleted again before storage at -80 ℃ until proteomics were performed.

### Analytical methods

#### Cell density measurement

Cell density was measured by absorbance at 600 nm in the Thermo Scientific GENESYS 30 Visible Spectrophotometer (#840-277000).

#### Proteomics sample preparation

Pelleted cells were collected and lysed using bead beating, SDS-based lysis, and denaturation, followed by reduction with dithiothreitol and cysteine alkylation with iodoacetamide, as previously described(55). The protein aggregation capture (PAC) method was used to capture proteins and digest them *in situ* with sequencing-grade trypsin at a 1:75 (w/w) enzyme-to-protein ratio, initially incubated overnight at 37 °C and subsequently for 4 hr at room temperature (55). Tryptic digests were then acidified with 0.5% formic acid and filtered them through 10 kDa polyethersulfone (PES) spin filters (Vivaspin columns; Sartorius). Peptide concentrations were measured using a NanoDrop spectrophotometer at an absorbance of 205 nm.

#### Liquid chromatography with tandem mass spectrometry (LS-MS/MS)

LC-MS/MS analysis was performed by directly injecting approximately 3 µg of each tryptic peptide sample onto an in-house pulled nanospray emitter (75 µm ID) packed with 15 cm of C18 resin (Kinetex, Phenomenex). Peptides were separated using a 180-min linear organic gradient on a Vanquish UHPLC system coupled directly to a nanoelectrospray source on a Q Exactive Plus mass spectrometer (Thermo Scientific). We measured and sequenced eluting peptides using the data-dependent acquisition, as previously described(28, 29, 55-57).

### Bioinformatics and data analysis

#### Phage genome annotation

The base annotation used was RefSeq Accession NC_008583.1. For proteins with putative or hypothetical functions, homologs in well-studied staphylococcal phages *ϕ*11, *ϕ*NM2, and 80⍺ were identified based on synteny and nucleotide similarity from a progressiveMauve alignment(58). Protein homologs were confirmed via BLASTp with default settings, and the homologous function was assigned to the *ϕ*NM1 protein based on a literature search. If this approach was unsuccessful in annotating a protein, two searches were performed: 1.) a BLASTp homology search with default settings against the ClusteredNR database, and 2.) a Foldseek structural search with default settings against all available databases(59). Proteins that were still unable to be annotated after this pipeline were considered to have unknown functions.

#### Growth kinetics

Cell growth was determined for pre-, mid-, and late-infection periods, assuming first-order kinetics. Specific growth rate is determined as follows:

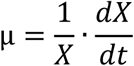

where μ (h^-1^) is the specific growth rate, X (g/L) is the cell concentration, which is proportional to OD_600nm_, and t (h) is the culturing time.

#### Proteomics

Collected MS/MS spectra were analyzed against sample-specific proteomic databases for *S. aureus*, *ϕ*NM1, and pDD0001 using the SEQUEST HT algorithm in Proteome Discoverer v2.3 (Thermo Scientific). First, peptide spectrum matches (PSMs) were scored and filtered using Percolator, setting the false discovery rate (FDR) to be <1% at both the PSM and peptide levels, as described previously(55). Next, protein-level abundance was quantified by calculating chromatographic area under the curve for each peptide, mapping them to their corresponding proteins, and summing their intensities. For downstream analysis, protein abundances were log_2_-transformed, LOESS normalized across replicates, and median-centered across all samples. For missing values, imputation was used to approximate the lower limit of detection of the mass spectrometer instrument. Statistical analyses were performed to identify significant differences in protein abundance between relevant sample groups after FDR correction. All raw mass spectra was deposited in the MassIVE (accession number: MSV000099116) and ProteomeXchange (accession number: PXD068231) databases.

#### Differentially expressed protein (DEP) analysis

LOESS-normalized, imputed, and median-centered protein abundances were analyzed using the DEP R package v1.24.0(60). Significant proteins were filtered out based on a 5% FDR and a minimum log_2_ fold change of 1.

#### Protein co-expression analysis

Co-expression analysis was performed on log_2_-transformed, LOESS-normalized, imputed, and median-centered protein abundances using the corAndPvalue function with default settings from the WGCNA R package v1.72-5(61). The co-expression network plot was generated using the circlize R package v 0.4.16(62).

#### Proteome and amino acid allocation

We calculated the proteome allocation for a protein (*f*_*P*_*i*__) or a group of proteins (*f*_*COG*_*i*__) based on their relative fractions by dividing their raw protein abundances by the sum of all protein abundances(28, 29, 63). We used COGclassifier v1.0.5 to classify proteins into COGs(64). We determined amino acid allocation (*f*_*AA*_*i*__) as the normalized fractions of each amino acid in the measured proteomes, as previously described(28, 29, 63). Surface proteins SasA, SasF, SasG, Spa, and ClfA were unannotated by COGclassifier so were manually added to COG-V, Defense Mechanisms, because it was the most appropriate COG category.

#### ATP cost analysis

The ATP cost of each amino acid (*C*_*AA*_*i*__) was calculated using pathway information from the Tier 2 Curated Organism Database of *Staphylococcus aureus aureus* NCTC 8325 hosted on BioCyc(65), as previously described(66) (**Supplementary Table S3**). The differential cost of each amino acid in ATP equivalents per average was calculated as follows:

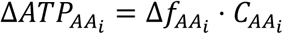

where 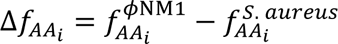 and the total cost difference in ATP per unit proteome between *ϕ*NM1 and *S. aureus* is 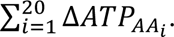

To calculate the cellular ATP requirements, the number of proteins per cell was approximated. First, the dry cell weight was calculated as follows:

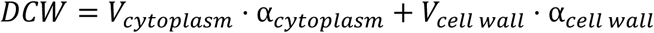

where *V*_*cytoplasm*_ and *V*_*cell wall*_ are the volumes of the cytoplasm and cell wall, respectively, and ⍺_*cytoplasm*_ and ⍺_*cell wall*_ are the dry weight densities of the cytoplasm and cell wall, respectively. The volumes are calculated according to the following equations:

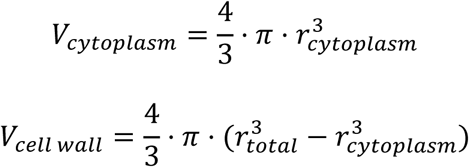

where *r*_*total*_ and *r*_*cytoplasm*_ are the radii of the total cell and cytoplasmic space, respectively, and *r*_*cytoplasm*_ = *r*_*total*_ − *t*_*cell wall*_ with *t*_*cell wall*_ being the thickness of the cell wall. Using parameter estimates of *t*_*cell wall*_ = 86 *nm*, *r*_*total*_ = 477 *nm*, 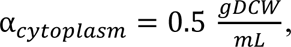 and 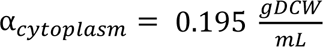 (67), *DCW* was determined to be 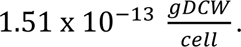 *MW_protein_*, the average molecular weight of a protein in *S. aureus*, was calculated as the unweighted average of the molecular weights of all measured proteins and was determined to be 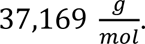 Assuming 55% of dry cell weight is protein, the number of proteins per cell was calculated as follows:

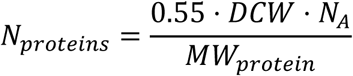

where *N*_*A*_ is Avogadro’s number. The total number of proteins per cell was estimated be 1.35 x 10^6^, which is consistent with previous estimates for *S. aureus* (**Supplementary File S1**) (68). The total number of ATP per cell required for synthesizing the proteome was calculated as follows:

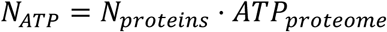

where *ATP*_*proteome*_ is the average protein synthesis cost for the proteome and is given by 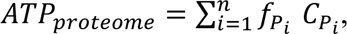 where *C*_*P*_*i*__ is the ATP cost for synthesizing protein i, calculated by

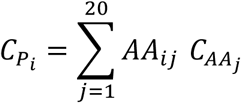

with *AA*_*ij*_ being the count of amino acid j in protein i. Using this methodology, it was determined that the average cost for synthesizing the *S. aureus* proteome was 1.8 x 10^9^ ATP/cell, which is reasonable compared to previous estimates in *E. coli* of 3.4 x 10^9^ ATP/cell(69) given that *S. aureus* has slightly less than half the cell volume as *E. coli* (∼0.45 µm^3^ vs. ∼1 µm^3^, respectively).

#### Statistical analysis

Statistical tests were performed using GraphPad Prism version 10.5.0 (GraphPad Software, Boston, Massachusetts, USA).

## Acknowledgements

This research was funded by the DARPA YFA award and Director Fellowship (D17AP00023) and DOE BER award (DE-SC0022226). The mention of trade names or commercial products in this article is solely for the purpose of providing scientific information and does not imply recommendation or endorsement by the funding agency.

## Supplementary Tables, Figures, Files, and Datasets

**Table S1:** Genome annotation of *ϕ*NM1.

**Table S2:** Strains, plasmids, phagemids, phages, and primers.

**Table S3:** Amino acid ATP costs.

**Figure S1:** Differential protein expression between lysogenic and non-lysogenic *S. aureus* strains before infection.

**Figure S2:** Cell growth kinetics of (**A**) non-lysogenic and (**B)** lysogenic *S. aureus* infected with *ϕ*NM1 or *ϕ*DD0001. Selected comparisons from two-way ANOVA are shown. Statistical analyses were performed in Graphpad Prism v10.6.0 with default settings.

**File S1:** Jupyter notebook for the calculation of the number of proteins per *S. aureus* cell.

**Dataset 1:** Protein allocation in non-lysogenic *S. aureus* infected by *ϕ*NM1 during pre-, mid-, and late-infection.

**Dataset 2:** Differentially expressed proteins in non-lysogenic *S. aureus* infected by *ϕ*NM1. Pairwise comparisons are presented for mid-infection versus pre-infection, late-infection versus pre-infection, and late-infection versus mid-infection.

**Dataset 3:** Protein allocation in lysogenic *S. aureus* infected by *ϕ*NM1 during pre-, mid-, and late-infection.

**Dataset 4:** Differentially expressed proteins in lysogenic and non-lysogenic *S. aureus* infected by *ϕ*NM1. Pairwise comparisons are presented for lysogen versus non-lysogen at pre-, mid-, and late-infection.

**Dataset 5:** Protein allocation in non-lysogenic *S. aureus* infected by *ϕ*DD0001 during pre-, mid-, and late-infection.

**Dataset 6:** Differentially expressed proteins in non-lysogenic *S. aureus* infected by *ϕ*DD0001 or *ϕ*NM1. Pairwise comparisons are presented for *ϕ*DD0001 versus *ϕ*NM1 at mid- and late-infection.

## References

1. M. R. Clokie, A. D. Millard, A. V. Letarov, S. Heaphy, Phages in nature. Bacteriophage 1, 31–45 (2011).

2. H. Ikeda, J.-i. Tomizawa (1968) Prophage P1, an extrachromosomal replication unit. in Cold Spring Harbor Symposia on Quantitative Biology (Cold Spring Harbor Laboratory Press), pp 791-798.

3. H. Brüssow, C. Canchaya, W.-D. Hardt, Phages and the evolution of bacterial pathogens: from genomic rearrangements to lysogenic conversion. Microbiology and molecular biology reviews 68, 560–602 (2004).

4. J. W. Little, Lysogeny, prophage induction, and lysogenic conversion. Phages: their role in bacterial pathogenesis and biotechnology, 37–54 (2005).

5. M. K. Waldor, J. J. Mekalanos, Lysogenic conversion by a filamentous phage encoding cholera toxin. Science 272, 1910–1914 (1996).

6. C. J. Murray et al., Global burden of bacterial antimicrobial resistance in 2019: a systematic analysis. The lancet 399, 629–655 (2022).

7. W. H. Organization, "Antimicrobial resistance global report on surveillance: 2014 summary" in Antimicrobial resistance global report on surveillance: 2014 summary. (2014).

8. T. Frieden, Antibiotic resistance threats in the United States. Centers Dis Control Prev 114 (2013).

9. S. A. Strathdee, G. F. Hatfull, V. K. Mutalik, R. T. Schooley, Phage therapy: From biological mechanisms to future directions. Cell 186, 17–31 (2023).

10. F. L. Gordillo Altamirano, J. J. Barr, Phage therapy in the postantibiotic era. Clinical microbiology reviews 32, 10.1128/cmr.00066-00018 (2019).

11. R. Capparelli, M. Parlato, G. Borriello, P. Salvatore, D. Iannelli, Experimental phage therapy against Staphylococcus aureus in mice. Antimicrobial agents and chemotherapy 51, 2765–2773 (2007).

12. B. A. Berryhill, D. L. Huseby, I. C. McCall, D. Hughes, B. R. Levin, Evaluating the potential efficacy and limitations of a phage for joint antibiotic and phage therapy of Staphylococcus aureus infections. Proceedings of the National Academy of Sciences 118, e2008007118 (2021).

13. M. Banar et al., A novel broad-spectrum bacteriophage cocktail against methicillin-resistant Staphylococcus aureus: Isolation, characterization, and therapeutic potential in a mastitis mouse model. Plos one 20, e0316157 (2025).

14. D. Bikard et al., Exploiting CRISPR-Cas nucleases to produce sequence-specific antimicrobials. Nature biotechnology 32, 1146–1150 (2014).

15. J. Y. Park et al., Genetic engineering of a temperate phage-based delivery system for CRISPR/Cas9 antimicrobials against Staphylococcus aureus. Scientific reports 7, 44929 (2017).

16. G. Ram, H. F. Ross, R. P. Novick, I. Rodriguez-Pagan, D. Jiang, Conversion of staphylococcal pathogenicity islands to CRISPR-carrying antibacterial agents that cure infections in mice. Nature biotechnology 36, 971–976 (2018).

17. L. H. Cobb et al., CRISPR-Cas9 modified bacteriophage for treatment of Staphylococcus aureus induced osteomyelitis and soft tissue infection. PLoS One 14, e0220421 (2019).

18. K. Kiga et al., Development of CRISPR-Cas13a-based antimicrobials capable of sequence-specific killing of target bacteria. Nature communications 11, 2934 (2020).

19. F.-Y. Li et al., Phagemid-based capsid system for CRISPR-Cas13a antimicrobials targeting methicillin-resistant Staphylococcus aureus. Communications Biology 7, 1129 (2024).

20. A. Brady et al., Molecular basis of lysis–lysogeny decisions in Gram-positive phages. Annual review of microbiology 75, 563–581 (2021).

21. T. B. Jacobson, M. M. Callaghan, D. Amador-Noguez, Hostile takeover: how viruses reprogram prokaryotic metabolism. Annual Review of Microbiology 75, 515–539 (2021).

22. C. Howard-Varona et al., Phage-specific metabolic reprogramming of virocells. The ISME journal 14, 881–895 (2020).

23. R. Pantůček et al., Identification of bacteriophage types and their carriage in Staphylococcus aureus. Archives of virology 149, 1689–1703 (2004).

24. C. Goerke et al., Diversity of prophages in dominant Staphylococcus aureus clonal lineages. Journal of bacteriology 191, 3462–3468 (2009).

25. M. Dini, L. Shokoohizadeh, F. A. Jalilian, A. Moradi, M. R. Arabestani, Genotyping and characterization of prophage patterns in clinical isolates of Staphylococcus aureus. BMC research notes 12, 1–6 (2019).

26. A. Ene et al., Examination of Staphylococcus aureus prophages circulating in Egypt. Viruses 13, 337 (2021).

27. A. Mayorga-Ramos, J. Zúñiga-Miranda, S. E. Carrera-Pacheco, C. Barba-Ostria, L. P. Guamán, CRISPR-Cas-based antimicrobials: design, challenges, and bacterial mechanisms of resistance. ACS infectious diseases 9, 1283–1302 (2023).

28. H. Seo, R. J. Giannone, Y.-H. Yang, C. T. Trinh, Proteome reallocation enables the selective de novo biosynthesis of non-linear, branched-chain acetate esters. Metabolic Engineering 73, 38–49 (2022).

29. C. Walker et al., Proteomes reveal metabolic capabilities of Yarrowia lipolytica for biological upcycling of polyethylene into high-value chemicals. Msystems 8, e00741–00723 (2023).

30. T. Shimada, Y. Yamazaki, K. Tanaka, A. Ishihama, The whole set of constitutive promoters recognized by RNA polymerase RpoD holoenzyme of Escherichia coli. PloS one 9, e90447 (2014).

31. D. Lindell et al., Genome-wide expression dynamics of a marine virus and host reveal features of co-evolution. Nature 449, 83–86 (2007).

32. K. Leskinen, B. G. Blasdel, R. Lavigne, M. Skurnik, RNA-sequencing reveals the progression of phage-host interactions between φR1-37 and Yersinia enterocolitica. Viruses 8, 111 (2016).

33. X. Zhao et al., Transcriptomic and metabolomics profiling of phage–host interactions between phage PaP1 and Pseudomonas aeruginosa. Frontiers in microbiology 8, 548 (2017).

34. X. Zhao et al., Global transcriptomic analysis of interactions between Pseudomonas aeruginosa and bacteriophage PaP3. Scientific reports 6, 19237 (2016).

35. R. Young, Bacteriophage lysis: mechanism and regulation. Microbiological reviews 56, 430–481 (1992).

36. T. J. Foster, Surface proteins of Staphylococcus aureus. Microbiology spectrum 7, 10.1128/microbiolspec.gpp1123-0046-2018 (2019).

37. F. S. Jacobson, R. Morgan, M. Christman, B. Ames, An alkyl hydroperoxide reductase from Salmonella typhimurium involved in the defense of DNA against oxidative damage: purification and properties. Journal of Biological Chemistry 264, 1488–1496 (1989).

38. G. Richarme, J. Abdallah, N. Mathas, V. Gautier, J. Dairou, Further characterization of the Maillard deglycase DJ-1 and its prokaryotic homologs, deglycase 1/Hsp31, deglycase 2/YhbO, and deglycase 3/YajL. Biochemical and biophysical research communications 503, 703–709 (2018).

39. C. Lee, C. Park, Bacterial responses to glyoxal and methylglyoxal: reactive electrophilic species. International journal of molecular sciences 18, 169 (2017).

40. T. G. Dong et al., Generation of reactive oxygen species by lethal attacks from competing microbes. Proceedings of the National Academy of Sciences 112, 2181–2186 (2015).

41. Y. Li et al., Oxidative stress elicited by phage infection induces Staphylococcal type III-A CRISPR–Cas system. Nucleic Acids Research 53, gkaf541 (2025).

42. V. Bohl et al., The Listeria monocytogenes persistence factor ClpL is a potent stand-alone disaggregase. Elife 12, RP92746 (2024).

43. C. Fang, Y. Zhang, Bacterial MerR family transcription regulators: activationby distortion: The mechanism of transcription regulation by MerR. Acta Biochimica et Biophysica Sinica 54, 25 (2021).

44. A. C. Eliot, J. F. Kirsch, Pyridoxal phosphate enzymes: mechanistic, structural, and evolutionary considerations. Annual review of biochemistry 73, 383–415 (2004).

45. P. Bilski, M. Li, M. Ehrenshaft, M. Daub, C. Chignell, Vitamin B6 (pyridoxine) and its derivatives are efficient singlet oxygen quenchers and potential fungal antioxidants. Photochemistry and photobiology 71, 129–134 (2000).

46. J. T. Rostøl, N. Quiles-Puchalt, P. Iturbe-Sanz, Í. Lasa, J. R. Penadés, Bacteriophages avoid autoimmunity from cognate immune systems as an intrinsic part of their life cycles. Nature Microbiology 9, 1312–1324 (2024).

47. J. Lossouarn et al., Enterococcus faecalis countermeasures defeat a virulent Picovirinae bacteriophage. Viruses 11, 48 (2019).

48. M. M. Susskind, D. Botstein, Mechanism of action of Salmonella phage P22 antirepressor. Journal of Molecular Biology 98, 413–424 (1975).

49. A. Das, S. Mandal, V. Hemmadi, V. Ratre, M. Biswas, Studies on the gene regulation involved in the lytic–lysogenic switch in Staphylococcus aureus temperate bacteriophage Phi11. The journal of biochemistry 168, 659–668 (2020).

50. M. M. Neamah et al., Sak and Sak4 recombinases are required for bacteriophage replication in Staphylococcus aureus. Nucleic Acids Research 45, 6507–6519 (2017).

51. B. J. Mendoza, X. Zheng, J. C. Clements, C. Cotter, C. T. Trinh, Potency of CRISPR-Cas Antifungals Is Enhanced by Cotargeting DNA Repair and Growth Regulatory Machinery at the Genetic Level. ACS Infectious Diseases 10.1021/acsinfecdis.3c00342 (2023).

52. W. Chen, Y. Zhang, W.-S. Yeo, T. Bae, Q. Ji, Rapid and efficient genome editing in Staphylococcus aureus by using an engineered CRISPR/Cas9 system. Journal of the American Chemical Society 139, 3790–3795 (2017).

53. D. G. Gibson et al., Enzymatic assembly of DNA molecules up to several hundred kilobases. Nature methods 6, 343–345 (2009).

54. O. Schneewind, D. Missiakas, Genetic manipulation of Staphylococcus aureus. Current protocols in microbiology 32, 9C. 3.1–9C. 3.19 (2014).

55. D. Dooley, et al., Expanded genome and proteome reallocation in a novel, robust Bacillus coagulans strain capable of utilizing pentose and hexose sugars. mSystems 9, e00952-00924 (2024).

56. C. Walker et al., Exploring Proteomes of Robust Yarrowia lipolytica Isolates Cultivated in Biomass Hydrolysate Reveals Key Processes Impacting Mixed Sugar Utilization, Lipid Accumulation, and Degradation. mSystems 6, e00443–00421 (2021).

57. C. Walker, S. Ryu, R. J. Giannone, S. Garcia, C. T. Trinh, Understanding and Eliminating the Detrimental Effect of Thiamine Deficiency on the Oleaginous Yeast *Yarrowia lipolytica*. Applied and Environmental Microbiology 86, e02299–02219 (2020).

58. A. E. Darling, B. Mau, N. T. Perna, progressiveMauve: multiple genome alignment with gene gain, loss and rearrangement. PloS one 5, e11147 (2010).

59. M. Van Kempen et al., Fast and accurate protein structure search with Foldseek. Nature biotechnology 42, 243–246 (2024).

60. X. Zhang et al., Proteome-wide identification of ubiquitin interactions using UbIA-MS. Nature protocols 13, 530–550 (2018).

61. P. Langfelder, S. Horvath, WGCNA: an R package for weighted correlation network analysis. BMC bioinformatics 9, 559 (2008).

62. Z. Gu, L. Gu, R. Eils, M. Schlesner, B. Brors, "Circlize" implements and enhances circular visualization in R. (2014).

63. D. Dooley, et al., Expanded genome and proteome reallocation in a novel, robust Bacillus coagulans strain capable of utilizing pentose and hexose sugars. MSystems 9, e00952-00924 (2024).

64. Y. Shimoyama (2022) COGclassifier: A tool for classifying prokaryote protein sequences into COG functional category.

65. P. D. Karp et al., The BioCyc collection of microbial genomes and metabolic pathways. Briefings in bioinformatics 20, 1085–1093 (2019).

66. C. Kaleta, S. Schäuble, U. Rinas, S. Schuster, Metabolic costs of amino acid and protein production in Escherichia coli. Biotechnology journal 8, 1105–1114 (2013).

67. P. J. Wyatt, Cell wall thickness, size distribution, refractive index ratio and dry weight content of living bacteria (stapylococcus aureus). Nature 226, 277–279 (1970).

68. D. Zühlke et al., Costs of life-Dynamics of the protein inventory of Staphylococcus aureus during anaerobiosis. Scientific reports 6, 28172 (2016).

69. W. F. Martin, ATP requirements for growth reveal the bioenergetic impact of mitochondrial symbiosis. Biochimica et Biophysica Acta (BBA)-Bioenergetics, 149564 (2025).

