## Supplemental Materials for "Proteome Reallocation Reveals Dynamics and Mechanisms of Phage Infection and Defense in *Staphylococcus aureus*"

**Table S1:** Genome annotation of $\phi$NM1.

| Uniprot ID | NCBI Locus Tag | NCBI Protein ID | Gene ID | Product | Re-annotated? | Method | Classifi-cation |
|---|---|---|---|---|---|---|---|
| A0EWH2 | SAPPV1_gp01 | YP_873950.1 | GeneID:4525429 | Integrase | N | Refseq | Known |
| A0EWH3 | SAPPV1_gp02 | YP_873951.1 | GeneID:4525430 | Ith-2 | Y | Homology | Known |
| A0EWH4 | SAPPV1_gp03 | YP_873952.1 | GeneID:4525431 | Tha-2 | Y | Homology | Known |
| A0EWH5 | SAPPV1_gp04 | YP_873953.1 | GeneID:4525432 | CI-like repressor | N | Refseq | Known |
| A0EWH6 | SAPPV1_gp05 | YP_873954.1 | GeneID:4525388 | Cro-like repressor | N | Refseq | Known |
| A0EWH9 | SAPPV1_gp08 | YP_873957.1 | GeneID:4525391 | Antirepressor | N | Refseq | Known |
| A0EWI2 | SAPPV1_gp11 | YP_873960.1 | GeneID:4525394 | Cell division inhibitor Gp11 | Y | Homology | Known |
| A0EWI4 | SAPPV1_gp13 | YP_873962.1 | GeneID:4525396 | SaPIbov2 derepressor | Y | Homology | Known |
| A0EWI5 | SAPPV1_gp14 | YP_873963.1 | GeneID:4525397 | Sak4-like recombinase | Y | Homology | Known |
| A0EWI6 | SAPPV1_gp15 | YP_873964.1 | GeneID:4525398 | Single strand DNA binding protein | N | Refseq | Known |
| A0EWI9 | SAPPV1_gp18 | YP_873967.1 | GeneID:4525401 | Replication initiation protein | N | Refseq | Known |
| A0EWJ0 | SAPPV1_gp19 | YP_873968.1 | GeneID:4525402 | DnaC-like helicase loader | N | Refseq | Known |
| A0EWJ2 | SAPPV1_gp21 | YP_873970.1 | GeneID:4525404 | Endodeoxyribonuclease RusA | N | Refseq | Known |
| A0EWJ4 | SAPPV1_gp23 | YP_873972.1 | GeneID:4525406 | Panton-Valentine leukocidin | N | Refseq | Known |
| A0EWJ5 | SAPPV1_gp24 | YP_873973.1 | GeneID:4525407 | Nucleotide kinase | N | Refseq | Known |
| A0EWK2 | SAPPV1_gp31 | YP_873980.1 | GeneID:4525439 | dUTPase | N | Refseq | Known |
| A0EWK5 | SAPPV1_gp34 | YP_873983.1 | GeneID:4525427 | RinB-like transcriptional activator | N | Refseq | Known |
| A0EWK7 | SAPPV1_gp36 | YP_873985.1 | GeneID:4525428 | Transcriptional activator RinA | N | Refseq | Known |
| A0EWK8 | SAPPV1_gp37 | YP_873986.1 | GeneID:4525443 | Phage terminase small subunit | N | Refseq | Known |
| A0EWK9 | SAPPV1_gp38 | YP_873987.1 | GeneID:4525444 | Terminase large subunit | N | Refseq | Known |
| A0EWL0 | SAPPV1_gp39 | YP_873988.1 | GeneID:4525445 | Portal protein | N | Refseq | Known |
| A0EWL1 | SAPPV1_gp40 | YP_873989.1 | GeneID:4525446 | Ejection protein | N | Refseq | Known |
| A0EWL3 | SAPPV1_gp42 | YP_873991.1 | GeneID:4525448 | Head scaffolding protein | N | Refseq | Known |
| A0EWL4 | SAPPV1_gp43 | YP_873992.1 | GeneID:4525449 | Major head protein | N | Refseq | Known |
| A0EWL6 | SAPPV1_gp45 | YP_873994.1 | GeneID:4525451 | Head-tail connector protein | N | Refseq | Known |
| A0EWL7 | SAPPV1_gp46 | YP_873995.1 | GeneID:4525408 | Head-tail joining protein | N | Refseq | Known |
| A0EWL8 | SAPPV1_gp47 | YP_873996.1 | GeneID:4525409 | Tail completion protein | N | Refseq | Known |
| A0EWL9 | SAPPV1_gp48 | YP_873997.1 | GeneID:4525410 | Tail terminator protein | N | Refseq | Known |
| A0EWM0 | SAPPV1_gp49 | YP_873998.1 | GeneID:4525411 | Major tail protein | N | Refseq | Known |
| A0EWM1 | SAPPV1_gp50 | YP_873999.1 | GeneID:4525412 | Tail chaperone protein | N | Refseq | Known |
| A0EWM2 | SAPPV1_gp51 | YP_874000.1 | GeneID:4525413 | Tail assembly chaperone | N | Refseq | Known |
| A0EWM3 | SAPPV1_gp52 | YP_874001.1 | GeneID:4525414 | Phage tape measure protein | N | Refseq | Known |
| A0EWM4 | SAPPV1_gp53 | YP_874002.1 | GeneID:4525415 | Distal tail protein Dit | N | Refseq | Known |
| A0EWM5 | SAPPV1_gp54 | YP_874003.1 | GeneID:4525416 | Tail-associated lysin | Y | Homology | Known |

| | | | | | | | |
|---|---|---|---|---|---|---|---|
| A0EWM6 | SAPPV1_gp55 | YP_874004.1 | GeneID:4525417 | Receptor binding protein | Y | Homology | Known |
| A0EWM7 | SAPPV1_gp56 | YP_874005.1 | GeneID:4525418 | Lower tail fiber protein | Y | Homology | Known |
| A0EWM8 | SAPPV1_gp57 | YP_874006.1 | GeneID:4525419 | Tail fiber protein | N | Refseq | Known |
| A0EWN1 | SAPPV1_gp60 | YP_874009.1 | GeneID:4525422 | Tail-associated cell-wall hydrolase | N | Refseq | Known |
| A0EWN2 | SAPPV1_gp61 | YP_874010.1 | GeneID:4525423 | Upper tail fiber protein | Y | Homology | Known |
| A0EWN4 | SAPPV1_gp63 | YP_874012.1 | GeneID:4525425 | Holin | N | Refseq | Known |
| A0EWN5 | SAPPV1_gp64 | YP_874013.1 | GeneID:4525426 | Endolysin | N | Refseq | Known |
| A0EWI0 | SAPPV1_gp09 | YP_873958.1 | GeneID:4525392 | Nitrile hydratase accessory protein??? | Y | Foldseek | Putative |
| A0EWI8 | SAPPV1_gp17 | YP_873966.1 | GeneID:4525400 | Abiα-like protein | Y | Homology | Putative |
| A0EWJ6 | SAPPV1_gp25 | YP_873974.1 | GeneID:4525433 | Virulence associated | N | Refseq | Putative |
| A0EWJ7 | SAPPV1_gp26 | YP_873975.1 | GeneID:4525434 | RBP anchoring protein??? | Y | Foldseek | Putative |
| A0EWJ8 | SAPPV1_gp27 | YP_873976.1 | GeneID:4525435 | 50/60/70S Ribosomal Protein L18A mimic??? | Y | Foldseek | Putative |
| A0EWJ9 | SAPPV1_gp28 | YP_873977.1 | GeneID:4525436 | YopX protein domain-containing protein | N | Refseq | Putative |
| A0EWK3 | SAPPV1_gp32 | YP_873981.1 | GeneID:4525440 | Transcriptional regulator | N | Refseq | Putative |
| A0EWK4 | SAPPV1_gp33 | YP_873982.1 | GeneID:4525441 | Tsi2 antitoxin-like protein | Y | Foldseek | Putative |
| A0EWL5 | SAPPV1_gp44 | YP_873993.1 | GeneID:4525450 | Arc-like repressor | N | Refseq | Putative |
| A0EWN3 | SAPPV1_gp62 | YP_874011.1 | GeneID:4525424 | Collagenase-like protein | Y | Foldseek | Putative |
| A0EWH7 | SAPPV1_gp06 | YP_873955.1 | GeneID:4525389 | Hypothetical protein | N | Refseq | Unknown |
| A0EWH8 | SAPPV1_gp07 | YP_873956.1 | GeneID:4525390 | Hypothetical protein | N | Refseq | Unknown |
| A0EWI1 | SAPPV1_gp10 | YP_873959.1 | GeneID:4525393 | Hypothetical protein | N | Refseq | Unknown |
| A0EWI3 | SAPPV1_gp12 | YP_873961.1 | GeneID:4525395 | Hypothetical protein | N | Refseq | Unknown |
| A0EWI7 | SAPPV1_gp16 | YP_873965.1 | GeneID:4525399 | Hypothetical protein | N | Refseq | Unknown |
| A0EWJ1 | SAPPV1_gp20 | YP_873969.1 | GeneID:4525403 | Hypothetical protein | N | Refseq | Unknown |
| A0EWJ3 | SAPPV1_gp22 | YP_873971.1 | GeneID:4525405 | Hypothetical protein | N | Refseq | Unknown |
| A0EWK0 | SAPPV1_gp29 | YP_873978.1 | GeneID:4525437 | Hypothetical protein | N | Refseq | Unknown |
| A0EWK1 | SAPPV1_gp30 | YP_873979.1 | GeneID:4525438 | Hypothetical protein | N | Refseq | Unknown |
| A0EWK6 | SAPPV1_gp35 | YP_873984.1 | GeneID:4525442 | Hypothetical protein | N | Refseq | Unknown |
| A0EWL2 | SAPPV1_gp41 | YP_873990.1 | GeneID:4525447 | Hypothetical protein | N | Refseq | Unknown |
| A0EWM9 | SAPPV1_gp58 | YP_874007.1 | GeneID:4525420 | Hypothetical protein | N | Refseq | Unknown |
| A0EWN0 | SAPPV1_gp59 | YP_874008.1 | GeneID:4525421 | Hypothetical protein | N | Refseq | Unknown |

**Table S2:** Strains, plasmids, phagemids, phages, and primers.

| Type | Name | Parent | Sequence | Description | Source |
|---|---|---|---|---|---|
| Strain | *S. aureus* RN4220 | N/A | N/A | prophage-deficient, restriction-deficient, agrA mutant | BEI (NR-45946) |
| Strain | *S. aureus* Newman | N/A | N/A | harbors prophages ϕNM1–4 | ATCC (25904) |
| Strain | *S. aureus* SaDD0001 | *S. aureus* RN4220 | N/A | single ϕNM1 lysogen | This study |
| Strain | *S. aureus* SaDD1001 | *S. aureus* SaDD0001 | N/A | single ϕNM1 lysogen harboring pDD0001 | This study |
| Plasmid | pCasSA | N/A | N/A | Base plasmid | Addgene #98211 |
| Phagemid | pDD0001 | pCasSA | N/A | Base phagemid | This study |
| Phage | ϕNM1 | *S. aureus* Newman | N/A | Temperate phage #1 of *S. aureus* Newman | ATCC (25904) |
| Phage | ϕDD0001 | *S. aureus* SaDD1001 | N/A | Mixture of WT ϕNM1 and ϕNM1 particles with pDD0001 | This study |
| Primer | SA.f | N/A | GAAGTCAAATAAATCGCTTGC | Forward for amplifying only *S. aureus* (no ϕNMx homology, *nuc* gene) | This study |
| Primer | SA.r | N/A | TGTTGTTTAGCTTTATTTTGTGC | Reverse for amplifying only *S. aureus* (no ϕNMx homology, *nuc* gene) | This study |
| Primer | NM1.f | N/A | ATGTTATAGCTAGCCTTCGG | Forward for ϕNM1 check amplicon | This study |
| Primer | NM1.r | N/A | TTAAATTTCATGAGACAATAAACG | Reverse for ϕNM1 check amplicon | This study |
| Primer | NM2.f | N/A | TATCAAGATTAGAAGCGGGC | Forward for ϕNM2 check amplicon | This study |
| Primer | NM2.r | N/A | TTACGCAAGTAAGTCCTCTG | Reverse for ϕNM2 check amplicon | This study |
| Primer | NM3.f | N/A | AACGACGTGATGGTAATAGG | Forward for ϕNM3 check amplicon | This study |
| Primer | NM3.r | N/A | GAAAGCTATGAGCGTAATGC | Reverse for ϕNM3 check amplicon | This study |
| Primer | NM4.f | N/A | ACAGGGGTATGACATTGTTC | Forward for ϕNM4 check amplicon | This study |
| Primer | NM4.r | N/A | AAACCGGTCATTCTCTAACG | Reverse for ϕNM4 check amplicon | This study |
| Primer | pCasSA.f | N/A | AATATTGGTGAGTACGATGACGAAAGTTAAGTCCAGAAGGTCGATAGAAAGC | Forward for pCasSA backbone for *pac* insertion | This study |
| Primer | pCasSA.r | N/A | TCGTTCATTTCATTTACCACCAACTCTCGCGGTATTTACCACAACAGTACGCC | Reverse for pCasSA backbone for *pac* insertion | This study |
| Primer | pac.f | N/A | GCGAGAGTTGGTGGTAAATG | Forward for *pac* site insert | This study |
| Primer | pac.r | N/A | TTAACTTTCGTCATCGTACTCACC | Reverse for *pac* site insert | This study |

**Table S3:** Amino acid ATP costs.

| Amino Acid | ATP Cost (mol/mol) |
|---|---|
| Ala | 3.2 |
| Arg | 6.2 |
| Asn | 6.2 |
| Asp | 4.2 |
| Cys | 2.2 |
| Gln | 0.2 |
| Glu | -0.8 |
| Gly | 2.2 |
| His | 7.2 |
| Ile | 11.2 |
| Leu | -4.8 |
| Lys | 9.2 |
| Met | 9.2 |
| Phe | 4.2 |
| Pro | 4.2 |
| Ser | 2.2 |
| Thr | 10.2 |
| Trp | 6.2 |
| Tyr | 2.2 |
| Val | 2.2 |

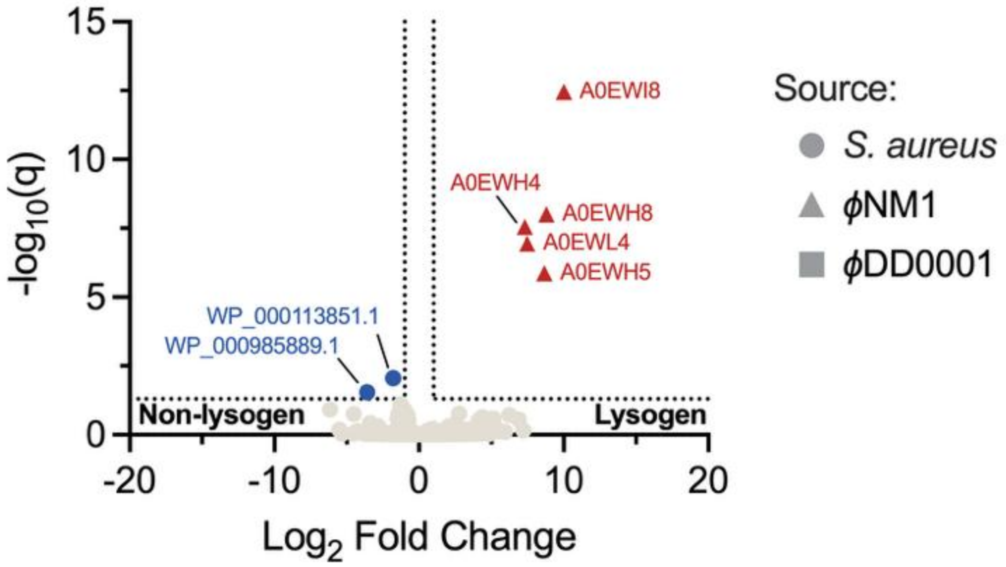


**Figure S1:** Differential protein expression between lysogenic and non-lysogenic *S. aureus* strains before infection.

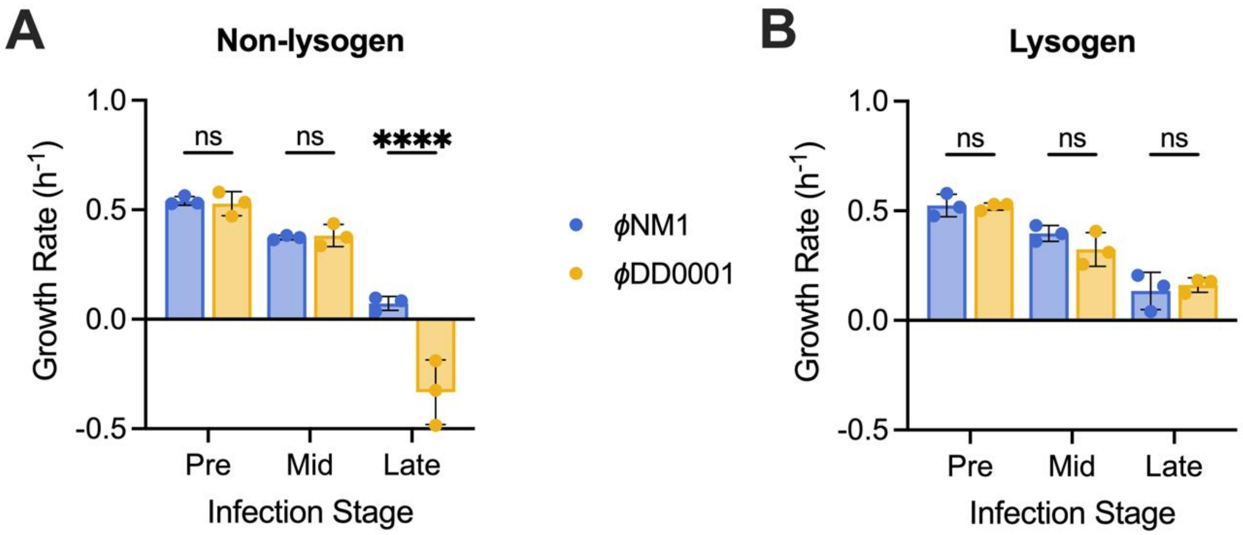


**Figure S2:** Cell growth kinetics of (**A**) non-lysogenic and (**B)** lysogenic *S. aureus* infected with $\phi$NM1 or $\phi$DD0001. Selected comparisons from two-way ANOVA are shown. Statistical analyses were performed in Graphpad Prism v10.6.0 with default settings.
